# Carbon-conserving Bioproduction of Malate in an *E. coli*-based Cell-Free System

**DOI:** 10.1101/2024.11.26.623433

**Authors:** Ryan A. L. Cardiff, Shaafique Chowdhury, Widianti Sugianto, Benjamin I. Tickman, Diego Alba Burbano, Pimphan A. Meyer, Margaret Cook, Brianne King, David Garenne, Alexander S. Beliaev, Vincent Noireaux, Pamela Peralta-Yahya, James M. Carothers

## Abstract

Formate, a biologically accessible form of CO_2_, has attracted interest as a renewable feedstock for bioproduction. However, approaches are needed to investigate efficient routes for biological formate assimilation due to its toxicity and limited utilization by microorganisms. Cell-free systems hold promise due to their potential for efficient use of carbon and energy sources and compatibility with diverse feedstocks. However, bioproduction using purified cell-free systems is limited by costly enzyme purification, whereas lysate-based systems must overcome loss of flux to background reactions in the cell extract. Here, we engineer an *E. coli*-based system for an eight-enzyme pathway from DNA and incorporate strategies to regenerate cofactors and minimize loss of flux through background reactions. We produce the industrial di-acid malate from glycine, bicarbonate, and formate by engineering the carbon-conserving reductive TCA and formate assimilation pathways. We show that *in situ* regeneration of NADH drives metabolic flux towards malate, improving titer by 15-fold. Background reactions can also be reduced 6-fold by diluting the lysate following expression and introducing chemical inhibitors of competing reactions. Together, these results establish a carbon-conserving, lysate-based cell-free platform for malate production, producing 64 μM malate after 8 hours. This system conserves 43% of carbon otherwise lost as CO_2_ and incorporates 0.13 mol CO_2_ equivalents/mol glycine fed. Finally, techno-economic analysis of cell-free malate production from formate revealed that the high cost of lysate is a key challenge to the economic feasibility of the process, even assuming efficient cofactor recycling. This work demonstrates the capabilities of cell-free expression systems for both the prototyping of carbon-conserving pathways and the sustainable bioproduction of platform chemicals.

**Highlights:**

- Successfully engineered the carbon-conserving reductive TCA and formate assimilation pathways in a lysate-based cell-free system for production of the C4 industrial di-acid malate from C1 and C2 feedstocks.
- Achieved a 6-fold reduction in competition from the endogenous cell-free metabolism by blocking TCA activity using small-molecule inhibitors and lysate dilution.
- Increased accumulation of malate by 15-fold in a single-step reaction using cell-free expression of an enzymatic cofactor regeneration system.
- Techno-economic analysis identified routes for economically feasible production of malate from renewable feedstocks in a cell-free system by improving conversion efficiency and reducing lysate cost.

**Graphical Abstract:** 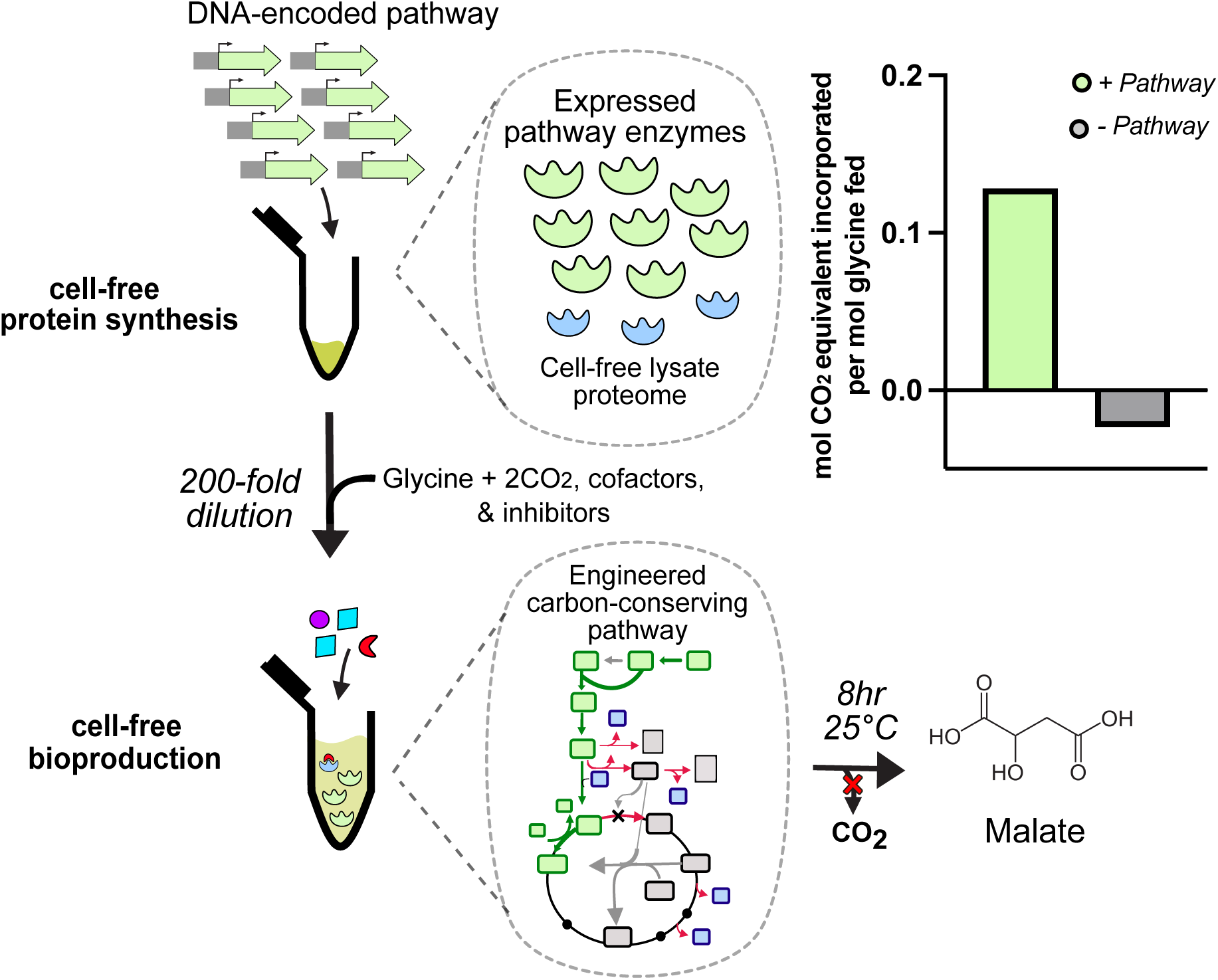

## Introduction

Metabolic engineering has enabled the production of industrially relevant chemicals from renewable and low-cost one carbon (C1) feedstocks, including CO_2_ and formate ^1–3^. The use of renewable feedstocks offers a promising alternative to petroleum-based chemical synthesis for the sustainable production of platform chemicals ^4,5^. However, the use of CO_2_ and formate as feedstocks for common metabolic engineering chassis, such as *Escherichia coli,* has been limited by the low solubility of CO_2_ and high formate toxicity ^6,7^. Novel approaches are needed to efficiently incorporate C1 feedstocks into biologically-accessible chemicals.

The use of cell-free gene expression systems (CFE) as a platform for carbon-conserving metabolic engineering has the potential to address several of the limitations of microbial based bioproduction ^8^. CFE are typically either prepared as a purified system, in which all the necessary components are individually purified and then reconstituted, or as a cell-based lysate ^9,10^. Each CFE approach has its own advantages and weaknesses, but across systems the high cost of preparation limits their potential for metabolic engineering ^11^. Lysate-based transcriptional and translational (TXTL) systems show promise for high-yield protein expression at lower costs than purified systems ^12^, while still enabling modification of reaction conditions, such as protein and chemical concentrations. CFE prepared from *E. coli* lysate contain high concentrations of cellular proteins and metabolites, including those required for transcriptional and translation, but lack the cell membrane and genomic DNA native to the cell ^13^. This allows CFE to be leveraged as an open reaction vessel, where plasmid or linear DNA encoding pathways of interest can be added directly to the reaction mixture.

For lysate-based metabolic engineering, there are many outstanding questions with respect to the relevant expression levels of multi-enzyme pathways, compatibility of reaction conditions for gene expression and bioproduction, and the challenges associated with the endogenous proteome of the lysate. To date, the majority of *in vitro* carbon-fixing pathways have been assembled using purified enzyme systems to avoid interference from native enzymes present in a lysate-based system ^14–16^. However, purified systems are limited by the high cost of biocatalyst preparation ^14,17–19^. While less costly to prepare than purified enzyme systems, lysate-based systems face challenges from the presence of native pathways and an inability to easily regulate endogenous enzyme activity in CFE ^20,21^. While the activity of the endogenous proteome in cell-free lysates is well-documented ^18,22^, effective interventions are still needed to minimize diversion of flux from competing reactions in the proteome. Developing systems to understand which enzymes and metabolic branches are important focal points for diversion is crucial for cell-free metabolic engineering and bioproduction more broadly.

Several carbon-fixing or carbon-conserving bioproduction pathways have been demonstrated using purified enzyme systems. Many systems take advantage of carboxylase-dependent cycles to iteratively fix carbon into a C3 or C4 product ^14–16^. Carboxylase-dependent pathways are often easier to implement *in vitro* as they tend to avoid the need for unstable, oxygen-sensitive cofactors. However, these cyclical pathways tend to be energetically expensive in terms of NAD(P)H and ATP consumption compared to linear pathways, such as the reverse glycine cleavage pathway. However, linear C1-incorporating pathways may require the use of complex cofactors for key enzymes, such as vitamin B_12_, ferredoxins, or rare metals ^23–25^. Across pathways, efficient regeneration of energy sources and cofactors is needed to improve reaction productivity and reduce costs for C1 assimilation pathways ^4,15,21^. Novel approaches to reduce biocatalyst requirements and regulate metabolic activity will expand the potential of cell-free systems as a scalable platform for metabolic engineering.

In this work, we demonstrate a platform for the carbon-conserving bioproduction of malate in a lysate-based CFE. Microbial bioproduction of the industrial di-acid malate requires re-routing flux through either the oxidative or reductive TCA (rTCA) cycle. The rTCA cycle represents the most carbon-efficient route, with a theoretical yield of 1 mol malate/mol pyruvate with an extra carbon coming from bicarbonate; compared to a theoretical yield of 0.5 mol malate/mol pyruvate through the oxidative TCA cycle, which features two decarboxylation steps ^26^. Microbial production through either pathway requires extensive genome engineering to introduce knockouts that will redirect metabolic flux towards malate accumulation. This often leads to cells that have significant growth defects, as the knocked out genes are typically involved in the TCA cycle or central carbon metabolism ^26–28^. Cell-free bioproduction has the potential to overcome these limitations by decoupling growth and bioproduction. Therefore, we sought to develop a cell-free platform for malate production through the carbon-efficient rTCA pathway.

The presented pathway incorporates two carbon-fixation steps to generate the C4 di-acid malate from the C2 precursor glycine and C1 inputs formate and bicarbonate (**Figure 1**). This system utilizes steps from both the formate assimilation pathway and reductive TCA cycle (rTCA). Recently, we have shown that the formate assimilation pathway coupled with the reductive glycine pathway (rGCV) can be used to assimilate formate into glycine and serine ^29^. This has shown to be the most energetically efficient of the natural C1 fixing pathways ^29^. In the formate assimilation pathway, formate is first coupled with tetrahydrofolate (THF) to ultimately generate the one-carbon donor, 5,10-methylenetetrahydrofolate (5,10-CH_2_-THF). Next, 5,10-CH_2_-THF transfers the C1 to glycine to synthesize serine and recycle THF. Serine is then converted to pyruvate, where another CO_2_ equivalent is incorporated through pyruvate carboxylase (*pyc*) to generate oxaloacetate. Oxaloacetate is then reduced to malate via malate dehydrogenase (*mdh*) using NADH. We find that the expression of formate dehydrogenase (*fdh*) to regenerate NADH helps drive metabolic flux towards malate by maintaining a high concentration of reducing equivalents.

**Figure 1.**
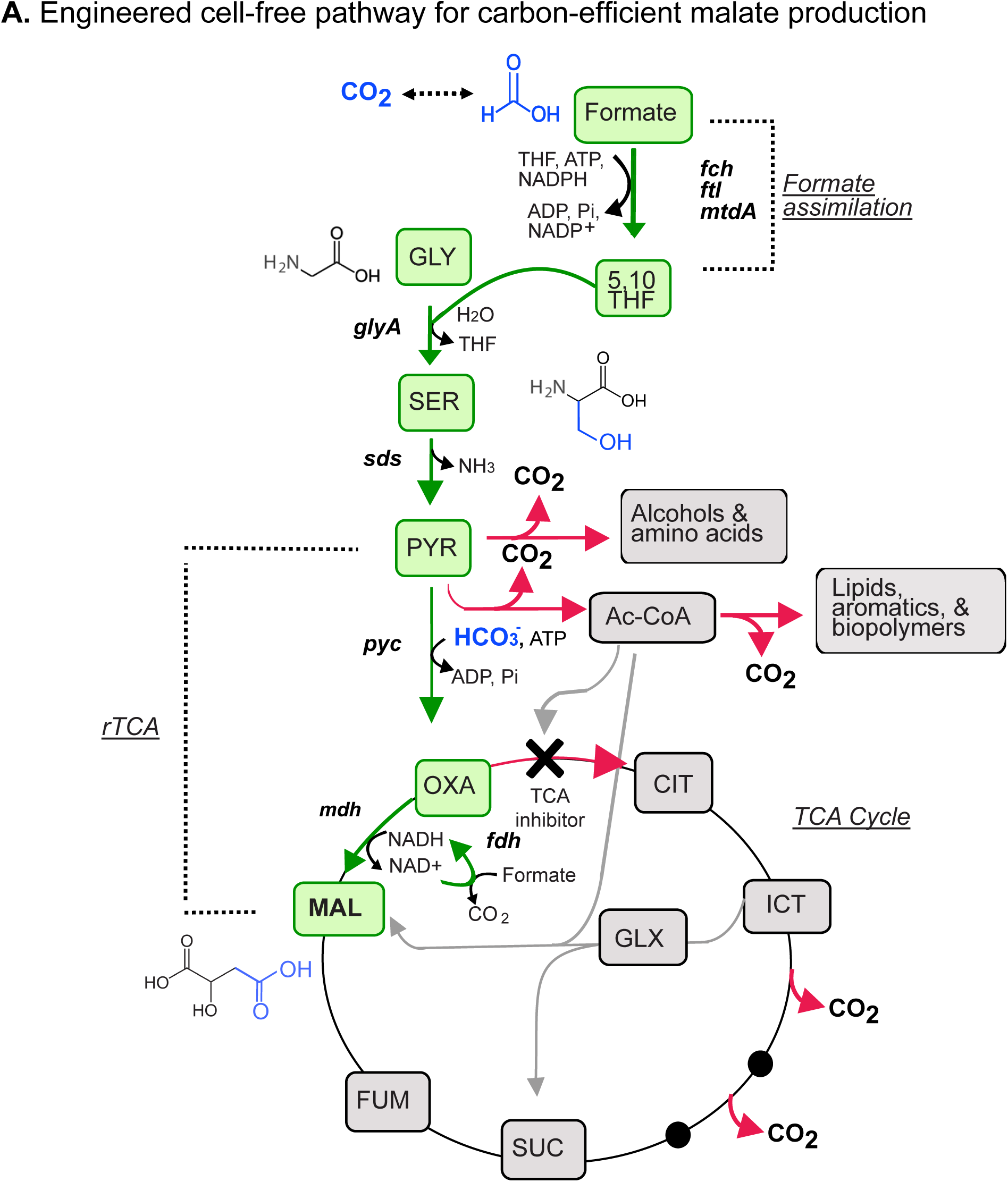
Overview of the *E. coli-*based cell-free system for malate production. A. Pathway diagram for conversion of formate and glycine into malate. Formate is incorporated into glycine to form serine via the four-step formate assimilation pathway. Serine is then converted to malate via the three-step reductive TCA cycle. High NADH concentrations are maintained by *in situ* regeneration from *fdh* to bias *mdh* flux towards malate. Gene abbreviations: *ftl*, formate-tetrahydrofolate ligase; *fch*, methenyltetrahydrofolate cyclohydrolase; *mtdA*, methylenetetrahydrofolate dehydrogenase (NADP^+^); *glyA*, glycine hydroxymethyltransferase; *sds,* serine dehydratase; *pyc*, pyruvate carboxylase; *mdh,* malate dehydrogenase; *fdh*, formate dehydrogenase. Metabolite abbreviations: THF, tetrahydrofolate; 5,10-THF, 5,10-methylene tetrahydrofolate; GLY, glycine; SER, serine; PYR, pyruvate; OXA, oxaloacetate; MAL, malate; SUC, succinate; ICT, isocitrate; CIT, citrate; GLX, glyoxylate; FUM,fumarate; Ac-CoA, acetyl-CoA.

Development of strategies to overcome limitations of lysate-based CFE metabolic engineering is crucial to the successful implementation of complex pathways. In TXTL systems, the optimal conditions for gene expression may not necessarily be optimal for biosynthesis. We separate the one-pot metabolic engineering platform into two parts by separating the cell-free protein synthesis reaction from the bioproduction reaction, allowing us to independently tune conditions for each process. By diluting protein synthesis reactions, we improve the volumetric efficiency of the bioproduction reaction and reduce the relative activity from endogenous enzymes in the lysate. We also show that metabolic flux from competing pathways can be blocked by the addition of small-molecule inhibitors directly to the bioproduction reaction. In addition to improving efficiency, this approach minimizes carbon loss through the TCA cycle without requiring engineering of the underlying strain. Together, these developments enable malate production from formate and glycine with a yield on glycine of 6.4% in an eight-hour biosynthesis reaction. This work helps establish the capabilities of CFE as a platform for combined protein synthesis and metabolic engineering, enabling carbon-conserving bioproduction of platform chemicals.

## Results

### Metabolomics analysis of the endogenous metabolic activity of the CFE

While the lysis and filtration steps in the preparation of CFE lysates remove bulky membrane proteins and genomic DNA, many native protein complexes are still present at high concentrations ^30^. We first sought to characterize the basal metabolic activity in the cell-free system by measuring the activity from endogenous enzymes in the system via targeted metabolomics.

To understand the relative carbon flux as a result of the endogenous enzymes present in the cell-free lysate, we spiked 1 mM of pathway intermediates into otherwise empty, plain cell-free reactions and measured the presence of relevant compounds, including malate, pyruvate, fumarate, glyoxylate, and succinate, over time (**Figure 2A**). We found that, relative to a plain reaction with only water added, CFE with 1 mM malate spiked in accumulated high concentrations of fumarate, glyoxylate, succinate, and pyruvate (**Figure 2B, 2C)**, indicating that TCA enzymes are highly active in the cell-free lysate. We found that 100% of malate added was rapidly converted to other TCA intermediates. In four hours, a molar ratio of roughly 1:1:4:4 (fumarate:succinate:glyoxylate:pyruvate) was achieved, with no detectable malate in the reaction. Additionally, there is a high interconversion between pyruvate and malate, as shown by the reactions spiked with either compound having significant flux towards the other. With the addition of 1mM glycine, serine concentrations stay roughly constant at 25 μM (**Figure 2C**). In contrast, spiking 1mM serine leads to a rapid accumulation of glycine, reaching 485 μM, or 48.5% conversion, after 8 hours. A thermodynamic analysis using Equilibrator indicated that glycine is more energetically favorable than serine (ΔG = +6.7 kJ/mol) ^31^, which may explain the larger flux towards glycine **(Figure S1)**. These results highlight the large effects of the endogenous proteome on metabolite levels in lysate-based cell-free systems. For efficient bioproduction, strategies will be needed to counteract these effects.

**Figure 2.**
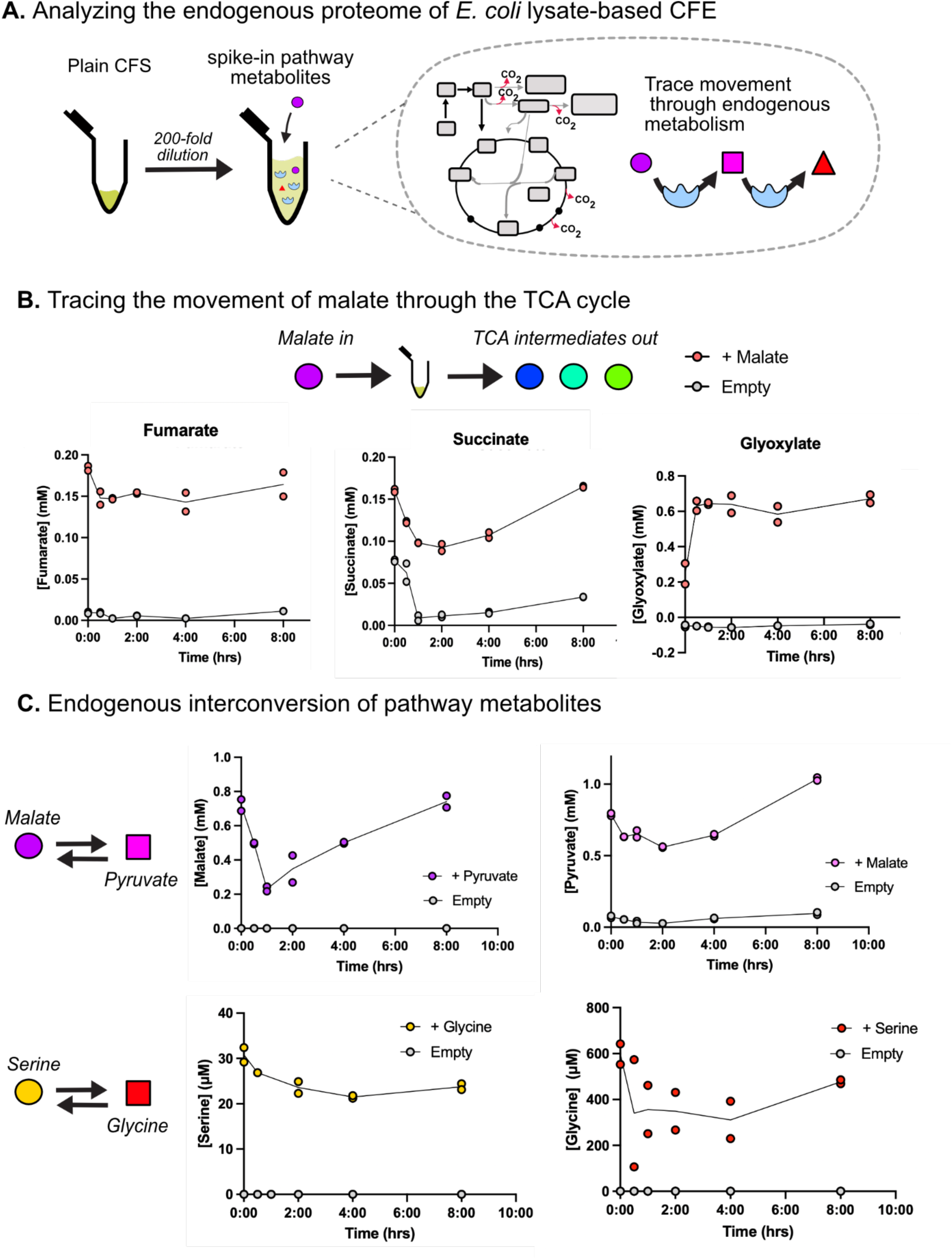
Analyzing the endogenous proteome of E. coli lysate-based CFE. A. Plain CFE reactions are set up with no genes expressed. Following overnight incubation, reactions are diluted 200-fold before introducing pathway metabolites. Metabolite concentrations are analyzed over eight hours. B. Flux through different TCA metabolites are traced over time for CFE reactions with or without malate added. Fumarate, succinate, and glyoxylate are shown as malate-derived metabolites with high flux. C. Interconversion of pathway metabolites is analyzed to understand the flux through endogenous enzymes in the system. For all panels, values represent the mean ± standard deviation of two technical replicates.

### Malate synthesis from the reductive TCA cycle

To engineer a carbon-efficient pathway for malate bioproduction, we began prototyping malate biosynthesis from the TCA cycle precursors oxaloacetate and pyruvate. We collected time-resolved measurements of malate production with cell-free expression of *E. coli* malate dehydrogenase (*mdh*) expressed from plasmid DNA with the p70 promoter. We found that in the case where *mdh* was overexpressed, malate accumulated rapidly, achieving over 100% conversion within 10 minutes **(Figure S2)**. Malate concentrations then rapidly decrease and reach baseline levels by 24 hours after the reaction start time. We hypothesized that malate flux was being diverted over time by other enzymes in the TCA cycle that consume malate or use NADH as a substrate ^32,33^. We then questioned if coupling malate synthesis with *in situ* NADH regeneration would allow us to continuously drive flux towards malate in our CFE. To favor *mdh* towards malate, we incorporated a NADH regeneration system based on a previously reported formate dehydrogenase (*fdh*) mutant from *Starkeya novella* ^34^. *fdh* can regenerate NADH and CO_2_ from formate and NAD^+^. Addition of NAD^+^ and formate to CFE with *fdh* expressed resulted in near complete conversion of NAD^+^ to NADH (**Figure 3A**). We then found that upon addition of NADH and excess formate to reactions that had co-expressed *mdh* and *fdh*, malate concentrations were stable at 1000 μM, representing full conversion efficiency, up to 24 hours (**Figure 3A**).

**Figure 3.**
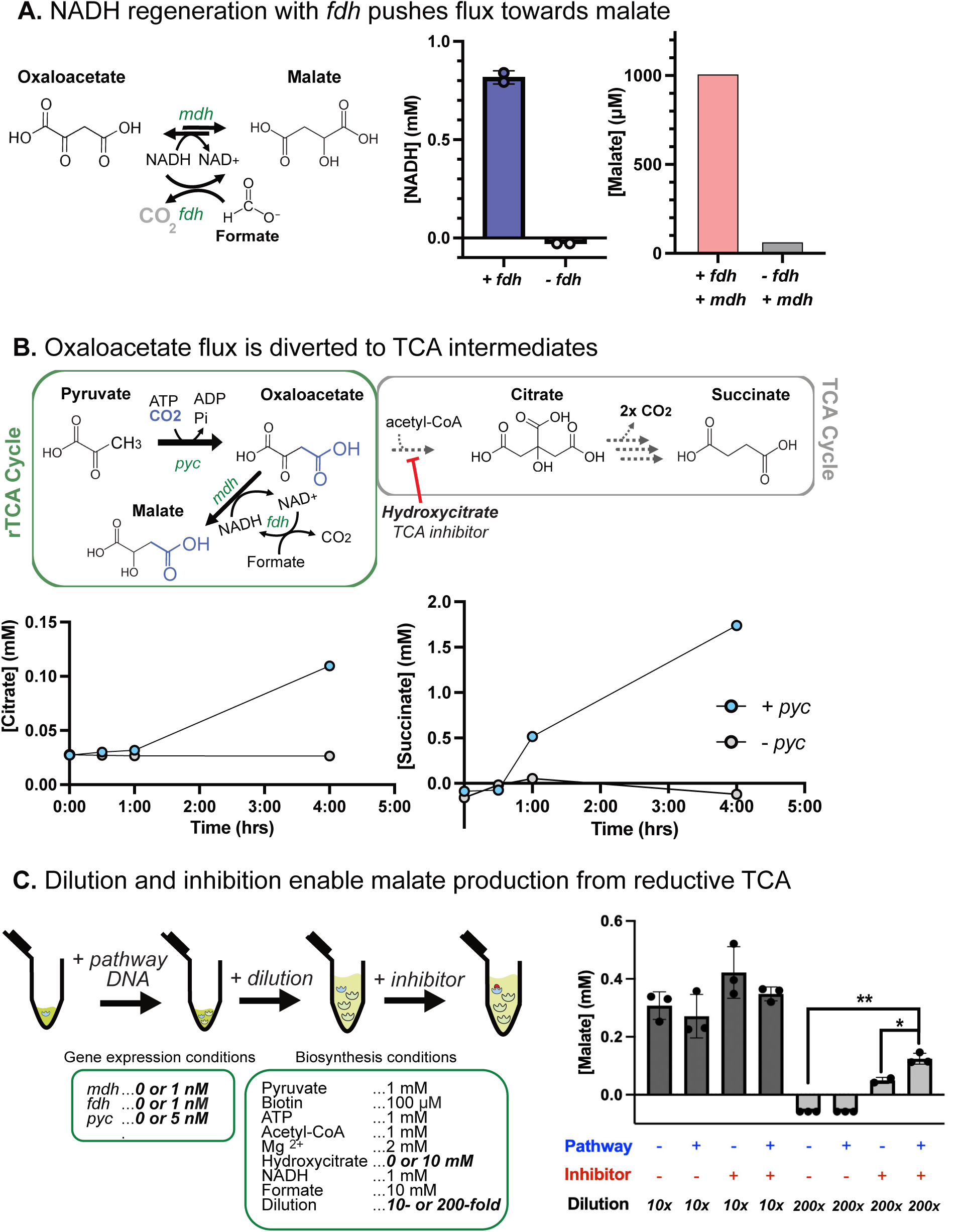
Optimization of the reductive TCA cycle. A. High malate levels are maintained by co-expression of *fdh* and *mdh*. Oxaloacetate is reduced to malate via *mdh* using NADH as the reducing equivalent. NADH can be regenerated from NAD^+^ and formate by *fdh* to maintain flux towards malate and prevent the reverse reaction of *mdh.* **Left:** NADH concentrations in reactions containing 1 mM NAD^+^, 10 mM formate, and CFE either with or without 5 nM *fdh* expression. **Right:** Malate concentrations in reactions containing 1 mM NADH, 1 mM oxaloacetate, and CFE with either 5 nM each of *mdh* and *fdh* or just *mdh*. B. *<u>pyc</u>* <u>expression results in diversion of oxaloacetate flux to TCA intermediates.</u> **Top:** Oxaloacetate and acetyl-CoA, an allosteric activator for *pyc*, can be fed into the TCA cycle via the irreversible citrate synthase (CS) enzyme. Entry into TCA can be inhibited by addition of the small molecule CS inhibitor, hydroxycitrate (HCT). Citrate **(bottom, left)** and succinate **(bottom, right)** concentrations are shown over time for reactions with or without pyruvate carboxylase (*pyc*) expressed. All reactions are diluted 200-fold for biosynthesis. Reactions contain 1 mM pyruvate, 10 mM HCO_3-_, 1 mM ATP, 2 mM Mg(CH₃COO)₂, 1 mM acetyl-CoA, 100 µM biotin, 1 mM NADH, and 10 mM formate. -*pyc* reactions express 1 nM of *mdh* and *fdh*, + *pyc* reactions also express 5 nM of *pyc*. C. CFE dilution and TCA inhibition enables carbon-efficient malate production from pyruvate. *pyc* incorporates one CO_2_ equivalent into pyruvate to form oxaloacetate and requires ATP, Mg^2+^, biotin, and acetyl-CoA for activation. Malate concentrations are measured after four hours for reactions with different conditions. DNA for pathway enzymes, additional CFE dilution, and the inhibitor hydroxycitrate are changed between conditions. All reactions contain 1 mM pyruvate, 10 mM HCO_3-_, 10 mM HCT, 1 mM acetyl-CoA, 100 µM biotin, 1 mM ATP, 2 mM Mg(CH_3_COO)_2_, 1 mM NADH, and 10 mM formate. Reactions contain CFE with or without 1 nM *mdh*, 1 nM *fdh*, and 5 nM *pyc*. Values represent the mean ± standard deviation of three technical replicates. For all panels, statistical significance was calculated using two-tailed unpaired Welch’s *t*-tests. Asterisks indicate a statistically significant difference (∗: p-value < 0.05, ∗∗: p-value < 0.005).

When *mdh* plasmid was omitted from the CFE, malate concentrations increased slowly over the reaction lifetime rather than accumulating quickly **(Figure S2)**. This is likely due to the presence of endogenous enzymes in the lysate that can natively produce malate. A proteomic analysis of *E. coli* cell-free lysate found that *mdh* and other TCA enzymes are among the most abundant enzymes in cell-free ^30^ **(Figure S3)**, supporting this hypothesis. Therefore, we questioned whether additional dilution of the cell-free reaction following protein expression could improve the pathway-dependent production of malate by minimizing the effects of competing endogenous enzymes. We tested three different dilutions of the CFE: 10 (our standard dilution factor), 50, and 200-fold. We found that the 200-fold dilution condition produced more malate than the other conditions when *mdh* was expressed **(Figure S4)**. Additionally, the 200-fold dilution produced less malate than the other dilutions when *mdh* was not expressed **(Figure S4)**, suggesting the activity of native *mdh* has also fallen below relevant levels in those conditions. These results may be due to a lower concentration of cofactors and substrates from the lysate, thereby reducing the relative activity of endogenous enzymes that convert malate into other products. Additionally, the dilution of the CFE may disrupt metabolon formation between TCA enzymes that accelerate malate loss ^35^.

Oxaloacetate, the direct C4 precursor of malate in our pathway, can be generated from C3 pyruvate and CO_2_ using the enzyme pyruvate carboxylase (*pyc*) from *Rhizobium etli*. The *pyc* enzyme is a large enzyme complex made up of three distinct domains: the ATP-consuming biotin carboxylase, carboxyltransferase, and biotin carboxyl carrier protein (BCCP) domain ^36^. We coupled the pyruvate carboxylase reaction with *mdh* and *fdh* to produce malate from the C3 precursor, pyruvate (**Figure 3B**). However, when we tested *pyc* activity from a 10-fold diluted CFE reaction, we observed no differential malate production in the conditions with and without *pyc*, with both conditions producing roughly 300 μM after four hours (**Figure 3C**, **S5)**. These results suggested that the engineered pathway may be outcompeted by the native carbon-catabolizing oxidative branch of the TCA cycle.

We hypothesized that oxaloacetate and pyruvate may be incorporated into the oxidative TCA cycle before being converted to malate by the heterologous pathway. Oxaloacetate and acetyl-CoA could be diverted into the endogenous oxidative TCA cycle by the enzyme citrate synthase, which is native to the lysate. The *pyc*-containing reactions are supplemented with acetyl-CoA, an allosteric activator shown to be essential for *pyc* activity ^37^. We hypothesized that greater dilutions of the CFE may improve flux through our engineered pathway, as pyruvate carboxylase has a lower reported dissociation constant for acetyl-CoA than that of citrate synthase ^36,38^. In the 10-fold diluted CFE condition, we did not see *pyc*-dependent malate production above the baseline levels generated by the endogenous lysate enzymes. In the 200-fold dilution condition, we observed increased *pyc*-dependent malate production compared to the baseline in the first hour of the reaction **(Figure S5)**, but after four hours the accumulated malate completely disappeared from both reactions (**Figure 3C**). Kinetic data also revealed a large *pyc-*dependent increase in the concentrations of the TCA cycle intermediates citrate and succinate (**Figure 3B**), consistent with a diversion of oxaloacetate flux into oxidative TCA by citrate synthase enzyme.

### Blocking TCA flux to improve pathway carbon-conservation

To minimize the loss of carbon through the oxidative TCA cycle, we investigated both targeted chemical inhibition of citrate synthase and re-routing of flux through overexpression of the glyoxylate shunt enzymes. Hydroxycitrate (HCT) is a potent competitive inhibitor of citrate synthase and ATP citrate lyase due to its structural similarity to citrate ^39^. We tested the ability of HCT to improve flux through our engineered pathway by adding varying amounts of hydroxycitrate to cell-free reactions and measuring malate accumulation over time. In the highest HCT concentration tested (10 mM), we see *pyc*-dependent accumulation of malate, reaching 212 ± 60 μM within four hours **(Figure S5)**. Additionally, we see accumulation of acetyl-CoA in the condition with the highest HCT concentration, implying that citrate synthase inhibition can prevent oxaloacetate consumption via the TCA cycle in cell-free reactions **(Figure S6)**. When combined, 200-fold dilution of the lysate and HCT inhibition lowered the background activity over 6-fold and improved pathway-dependent production 2.5-fold compared to 10-fold diluted reactions without TCA cycle inhibitor *(p < .05)* (**Figure 3C**) (**Table 1**).

**Table 1:**
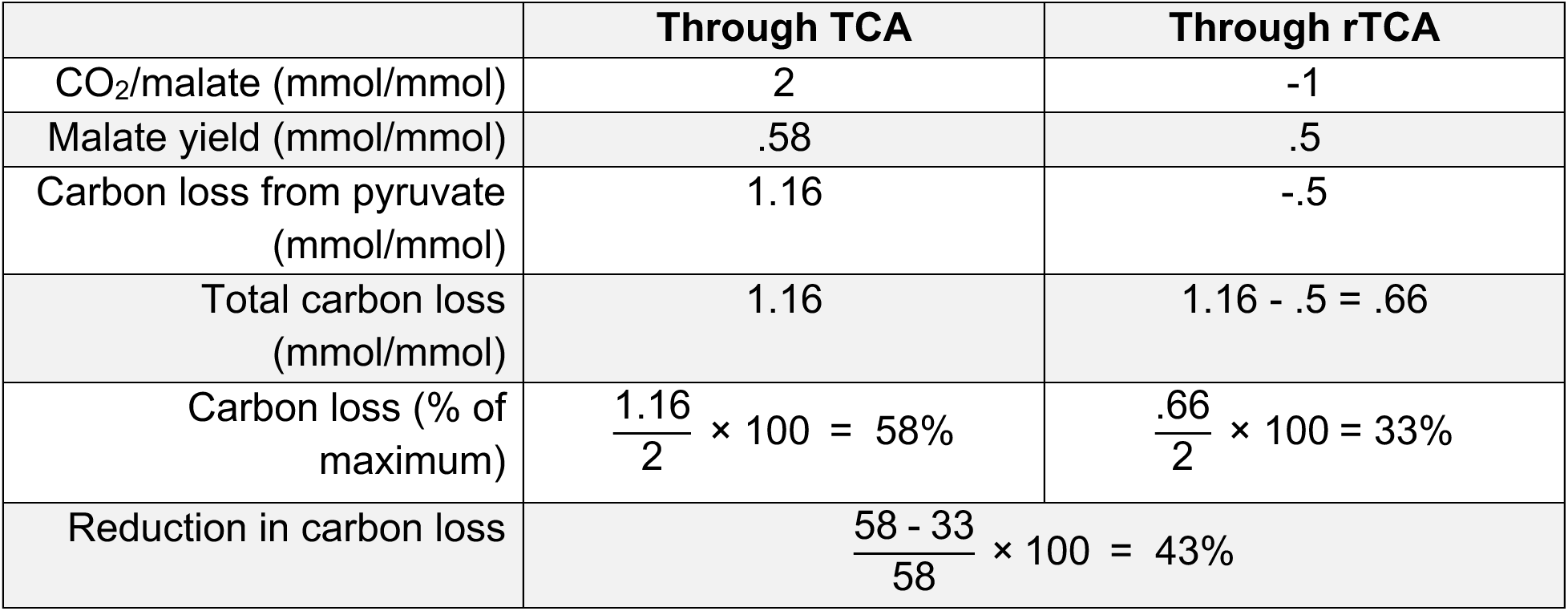
Carbon conservation for malate production through the rTCA pathway. Carbon conserved through the rTCA pathway is calculated for the production of malate from pyruvate. Data is shown in Figure 5B. The calculations assume a fixed stoichiometry for the moles of CO_2_ released per mole of malate generated through either the TCA or rTCA cycles.

As a second strategy for minimizing carbon loss, we tested whether overexpression of the glyoxylate shunt enzymes could redirect oxaloacetate flux that enters the TCA cycle towards malate. The glyoxylate shunt consists of two enzymes, isocitrate lyase (*ICL*) and malate synthase (*MS*), to convert isocitrate and acetyl-CoA to succinate and malate without loss of any carbon ^40^. We first confirmed that both enzymes generate measurable malate accumulation independently in a 200-fold diluted cell-free system **(Figure S7)** before combining them with the rest of the rTCA pathway. Surprisingly, when co-expressed with the rTCA pathway, we found malate titers decreased relative to the condition without the glyoxylate shunt **(Figure S7,** therefore, the glyoxylate shunt enzymes were not utilized in any of the studies described in the following sections. Understanding why the glyoxylate shunt was ineffective in these reactions would require further study.

### Balancing cofactor and expression requirements to construct multi-enzyme pathways

In the previous section, we showed that inhibiting citrate synthase effectively blocked TCA flux, allowing malate to be produced from pyruvate through the carbon-conserving rTCA pathway. To explore approaches for producing malate from simpler substrates, we aimed to extend our pathway to start from serine, which can be synthesized from the C2 substrate glycine and the C1 substrate formate via the rGCV pathway. Yu and Liao previously showed that serine can be efficiently converted into pyruvate through the heterologous expression of serine dehydratase (*sda*) in *E. coli* ^41^. We screened several isoforms of the *sda* enzyme from a variety of prokaryotic organisms in our standard CFE conditions and observed no catalytic activity (**Figure 4A**). As most prokaryotic isoforms of *sda* contain an oxygen-sensitive iron-sulfur cluster ^42^, we attempted to obtain enzyme function by supplementing iron and reducing agents, including dithiothreitol (DTT) and ascorbic acid at the time of expression. However, we still observed no clear activity from any of the *sda* isoforms screened (**Figure 4A**).

**Figure 4:**
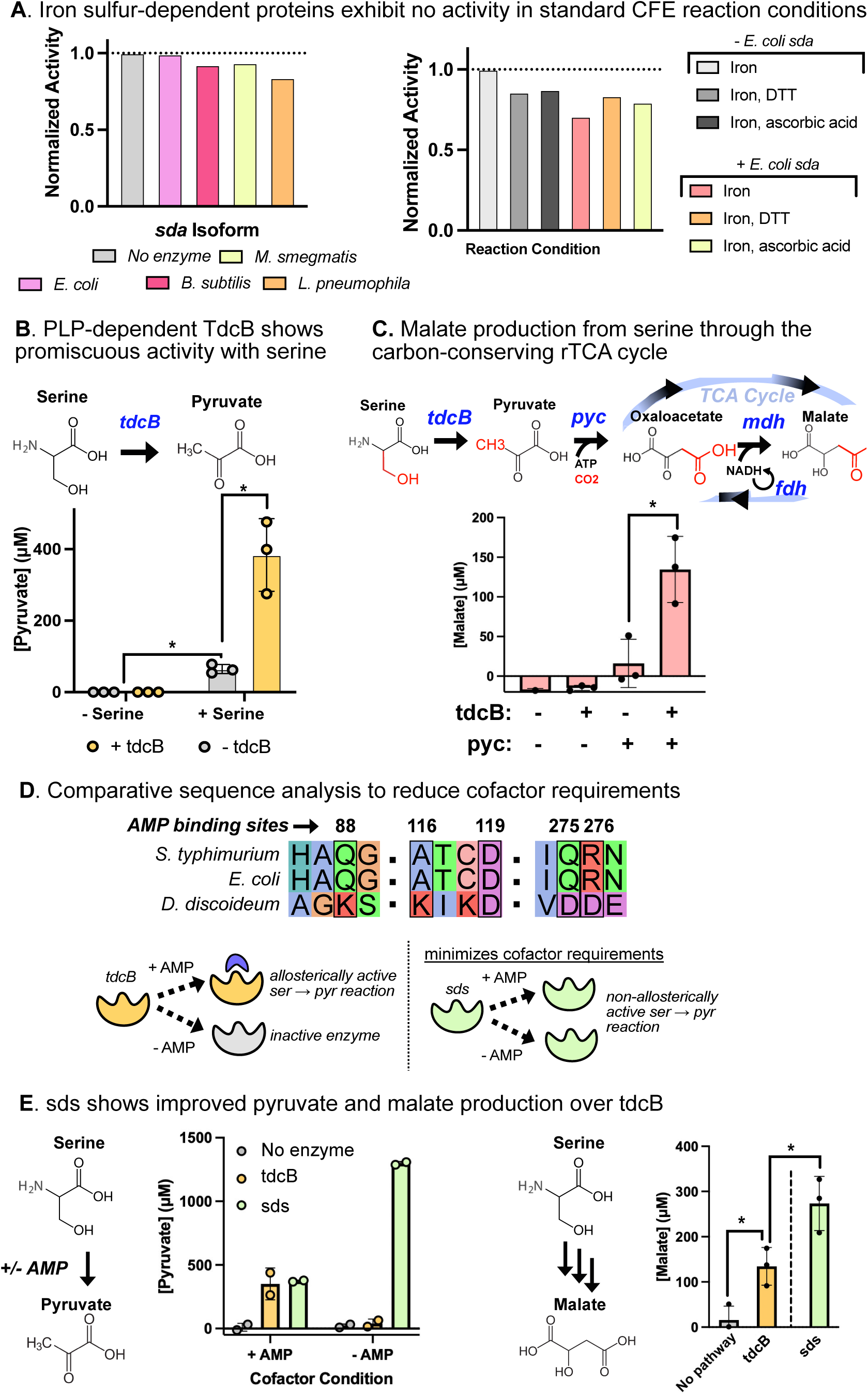
Characterization of serine deaminase function in CFE. A. Screening of iron-sulfur dependent serine deaminase (*sda*) variants in lysate-based CFE. **Left:** Isoforms from *E. coli, B. subtilis, M. smegmatis,* and *L. pneuomophila* were chosen from the literature based on favorable kinetic values and previous expression in *E. coli*. **Right:** Reaction conditions were screened to try to rescue activity of iron-sulfur enzymes in CFE. Iron was added to assist formation of the iron-sulfur cluster, and dithiothreitol or ascorbic acid were added to create a reducing environment. All genes were expressed at 5 nM and chemicals added at 500 µM. B. Pyruvate production is enabled using a promiscuous threonine deaminase (*tdcB)* from *E. coli. tdcB* belongs to the class of PLP-utilizing deaminases, avoiding the need for iron-sulfur proteins. Pyruvate concentrations are shown for reactions either with or without serine and *tdcB.* All reactions contain 100 µM PLP and 1 mM AMP. Values represent the mean ± standard deviation of three technical replicates. C. Serine is converted to malate using a four enzyme pathway in which *tdcB* converts serine to pyruvate, and pyruvate is carboxylated into oxaloacetate via *pyc*. Oxaloacetate is then reduced to malate by *mdh*, with cofactor regeneration from *fdh* to drive flux towards malate. Malate concentrations are shown for eight hour reactions containing combinations of *tdcB* and/or *pyc*, with all reactions containing *fdh* and *mdh*. D. Identification of an isoform via comparative sequence analysis removes the need for additional cofactors and reduces pathway incompatibilities. AMP binding sites found for the *tdcB* enzyme from *S. typhimurium* are compared against the *E. coli* isoform sequence and the *sds* sequence. E. Comparison of *sds* and *tdcB* for production of pyruvate (**left)** and malate (**right**). **Left:** Use of the *sds* isoform shows improved pyruvate production relative to *tdcB* in the absence of AMP. Pyruvate concentrations are shown for four hour reactions containing 1 mM serine, 100 µM PLP, and either with or without 1 mM AMP. Values represent the mean ± standard deviation of two technical replicates. **Right:** Comparison of *tdcB* and *sds* for serine to malate conversion via the four-enzyme pathway. Data shown for *tdcB* and *sds* are reproduced from Figures 4C and 5B, respectively. Values represent the mean ± standard deviation of three technical replicates. For all panels, statistical significance was calculated using two-tailed unpaired Welch’s *t*-tests. Asterisks indicate a statistically significant difference (∗: p-value < 0.05, ∗∗: p-value < 0.005).

We next tested the promiscuous serine to pyruvate conversion activity of a pyridoxal 5’-phosphate (PLP) dependent threonine dehydratase (*tdcB*) from *E. coli*. By adding exogenous serine into 200-fold diluted reactions, we were able to confirm serine-dependent *tdcB* activity, as *tdcB* expression produced 384 ± 101 μM pyruvate from 1 mM serine, compared to 65 ± 13 μM without *tdcB* expressed (**Figure 4B**). In the 10-fold diluted reaction condition, we did not observe differential production of pyruvate between the conditions with or without *tdcB* **(Figure S8)**. Based on these results and those in the previous section, we employed the 200-fold dilution condition for all subsequent reactions. Having demonstrated *tdcB* activity in a single-step transformation, we then integrated *tdcB* with the rTCA pathway to produce malate from serine. We measured malate production in four reaction conditions with and without *tdcB* and/or *pyc*, with all conditions containing *mdh* and *fdh*. At four hours, we saw no malate production across all conditions. However, we observed pyruvate accumulation in the conditions without *pyc* (**Figure S9)**, suggesting *pyc* was effectively pulling pyruvate flux forward. By eight hours, malate titers reached 135 μM when both *pyc* and *tdcB* were expressed, compared to no malate accumulation in the conditions without both enzymes expressed (**Figure 4C**).

In multi-enzyme cell-free systems, the use of enzymes with numerous cofactor requirements can reduce pathway efficiency and increase costs ^11,43^. Incorporation of ATP-regenerating systems in CFE can reduce cofactor requirements; however, we hypothesized that the addition of AMP as an allosteric activator for *tdcB* may interfere with ATP-dependent enzymes in the CFE ^9,44^. Therefore, we aimed to identify an alternative to *tdcB* that was insensitive to AMP regulation. From the literature, we extracted sequence motifs from *tdcB* known to interact with AMP ^45^. We then used multiple sequence alignment to choose other PLP-dependent enzymes that lacked these motifs and were predicted to be AMP-insensitive (**Figure 4D**, **Methods S1)** ^46^. From this screen, we selected a serine dehydratase (*sds*) variant from the soil-dwelling amoeba *Dictyostelium discoideum* to characterize in CFE ^47^. In the presence of AMP, the *sds* variant from *D. discoideum* produced similar levels of pyruvate as compared to *tdcB* from *E. coli* (**Figure 4E**). In reactions without AMP, *sds* achieved 100% conversion of serine to pyruvate after only four hours, while production from *tdcB* was indistinguishable from the background (**Figure 4E**). We then tested the ability of each isoform to convert serine to malate via the rTCA cycle. Here, we found that *sds* resulted in 2-fold higher malate concentrations compared to *tdcB* in an eight hour reaction (**Figure 4E**). Based on these results, subsequent pathway engineering was done with *sds* in place of *tdcB*.

### Connecting the reductive TCA cycle to formate assimilation products

In a fully integrated system for carbon-conserving malate production, the combined formate assimilation, rGCV, and rTCA pathways would fix 2 CO_2_ equivalents from formate and 2 from bicarbonate per malate produced. Glycine, the C2 product of the combined formate assimilation and rGCV pathways, can be converted to serine via the enzyme serine hydroxymethyltransferase (SHMT) encoded by the *glyA* gene from *E. coli*. *glyA* utilizes 5,10-CH_2_-THF as a C1 donor to assimilate a second formate-derived carbon into glycine (**Figure 1**). However, creating conditions for the rGCV pathway to run efficiently in the direction of glycine synthesis requires significant optimization of gene ratios and reaction tuning ^48–52^. In this section, we prototype *glyA* activity and estimate the efficiency of carbon incorporation into malate from glycine and 2 CO_2_ equivalents, 1 from formate and 1 from bicarbonate.

When fed with equimolar glycine and 5,10-CH_2_-THF, *glyA* converted 27% of the glycine into serine in four hours (269 ± 41 μM) in a 200-fold diluted cell-free reaction (**Figure 5A**). By incorporating *glyA* and *sds* with the rTCA genes, pathway-dependent malate production was observed directly from C2 glycine (**Figure 5B**, **top)**. Individual starting metabolites (pyruvate, serine, and glycine), were fed to batch reactions to analyze the efficiency of malate production from each step of the pathway. From pyruvate, we saw 1075 ± 36 μM malate in the presence of the pathway and 578 ± 113 μM with no pathway (**Figure 5B**, **left)**. When fed serine, the pathway produced 11-fold higher malate concentrations compared to the no-pathway control (273 ± 60 μM vs. 24 ± 8 μM) (**Figure 5B**, **left)**, suggesting the basal conversion of serine to pyruvate is low in CFE. From glycine, we saw 117 ± 6 μM malate produced with-and 62 ± 3 μM without-the pathway (**Figure 5B**, **left).**

**Figure 5.**
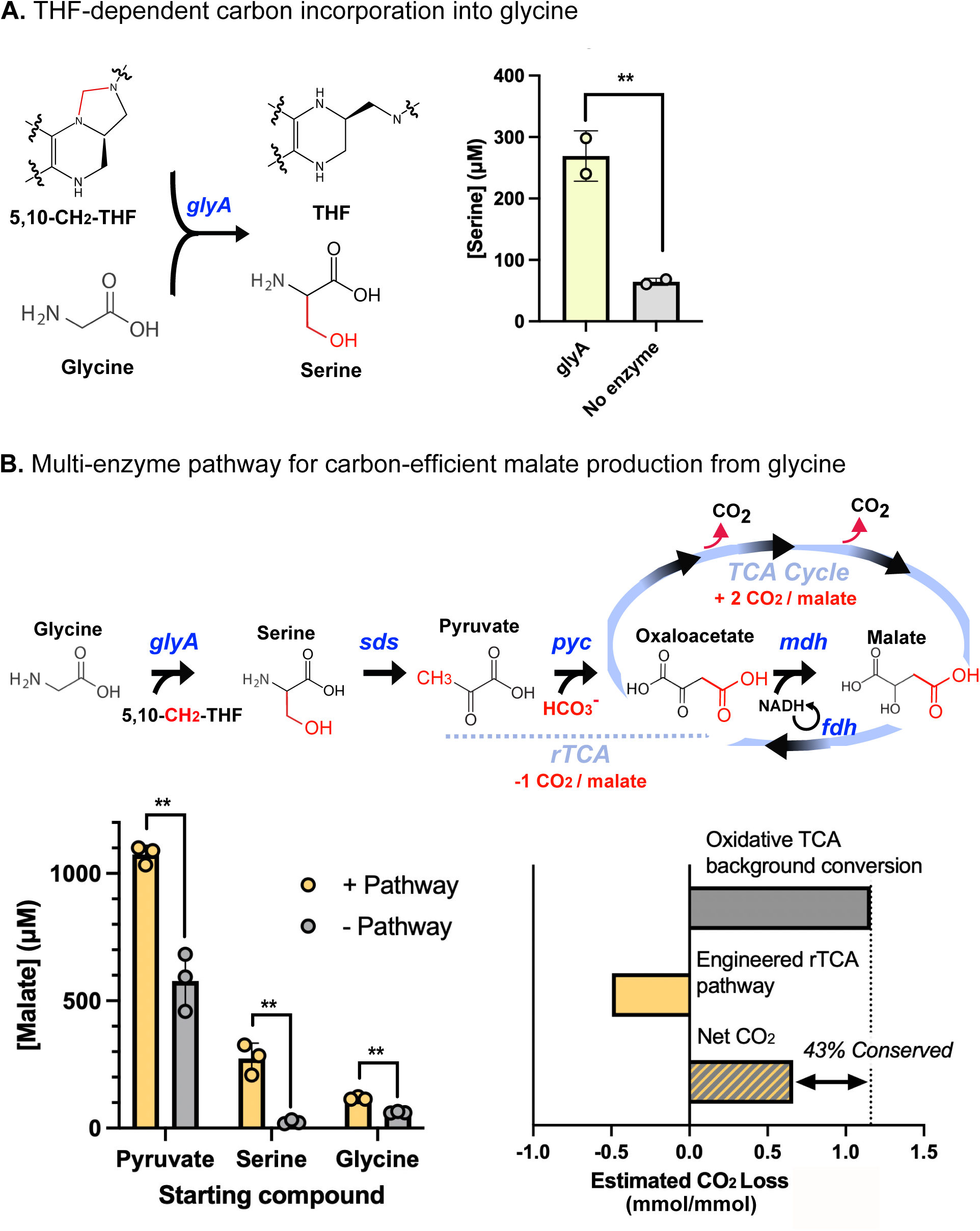
Conversion of glycine to malate. A. C1-incorporation into glycine, mediated by a THF intermediate. 5,10-methylenetetrahydrofolate (5,10-CH_2_-THF) acts as a C1 donor to convert glycine and water into serine, transforming 5,10-CH_2_-THF back to THF in the process. This process is catalyzed by the PLP-dependent *glyA* enzyme. Serine concentrations are shown for reactions containing 1 mM glycine, 1 mM 5,10-CH_2_-THF, <u>100 µM</u> PLP in conditions with and without 5 nM *glyA* expression. Values represent the mean ± standard deviation of two technical replicates. B. Multi-enzyme pathway enables carbon-efficient malate production from glycine in CFE. The pathway consists of five enzymatic steps, including *in situ* NADH regeneration. Incorporated carbons are shown in the schematic in red. **Left:** Malate concentrations are shown for multi-step reactions with different starting compounds either with or without the pathway expressed. **Right:** CO_2_ conserved through the introduction of the rTCA pathway is shown for pyruvate to malate conversion. CO_2_ conserved is estimated using the pathway-dependent conversion and assuming a fixed reaction stoichiometry between malate produced and CO_2_ equivalents conserved. Pathway-dependent conversion represents the added conversion efficiency above the background level from the pathway enzymes. Our calculations assume that for each pyruvate converted to malate, a maximum of 1 CO_2_ equivalent can be fixed from bicarbonate via rTCA, and a maximum of 2 CO_2_ equivalents can be released via the oxidative TCA cycle. For both panels, reactions contain all necessary chemical cofactors for the five-step pathway. Values represent the mean ± standard deviation of three technical replicates. For all panels, statistical significance was calculated using two-tailed unpaired Welch’s *t*-tests. Asterisks indicate a statistically significant difference (∗: p-value < 0.05, ∗∗: p-value < 0.005).

We estimated the amount of CO_2_ conserved in the engineered system using the pathway-dependent yield from the reactions fed with pyruvate. To calculate pathway-dependent yield, we took the difference in malate production with and without the engineered pathway. Then, to estimate conserved CO_2_, we assumed a fixed reaction stoichiometry between malate produced and CO_2_ equivalents conserved. Specifically, our calculations specify that for each pyruvate converted to malate, a maximum of 1 CO_2_ equivalent can be fixed from bicarbonate via rTCA, and a maximum of 2 CO_2_ equivalents can be released via the oxidative TCA cycle (**Table 1**).

Given the pathway-dependent differences in malate titers measured above, we estimate that the engineered pathway conserved 497 μM of CO_2_ compared to the oxidative TCA cycle when feeding pyruvate. This represents 50% of the potential conservation, assuming all of the malate accumulated above basal levels is produced through the engineered rTCA cycle (**Table 1**). The net result is that the introduction of the rTCA pathway increased malate yields by roughly two-fold (1075 vs. 578 μM malate) while reducing the estimated carbon loss as CO_2_ from the oxidative TCA cycle by 43% (0.66 vs. 1.2 mM CO_2_) (**Figure 5B, right) (Table 1**). Because the basal conversion of serine and glycine to malate from the endogenous lysate metabolism is low (**Figure 5B**, **left)**, we expect that the reduction in carbon loss remains similar when extending the engineered pathway to start from intermediates upstream from pyruvate.

### Prototyping cofactor regeneration to enable malate synthesis from formate

To construct the complete pathway for the conversion of formate and glycine to malate, 7 cofactors are required (**Figure 1**), 4 of which are consumed during biosynthesis (ATP, NADPH, NADH, and THF). In the previous section, we showed that glycine and 5,10-CH_2_-THF could be used to generate malate. In this section, we prototype the formate assimilation pathway for 5,10-CH_2_-THF regeneration to produce malate from glycine and formate directly. 5,10-CH_2_-THF can be generated from THF and formate in three enzymatic steps catalyzed by the ATP-dependent formate-THF ligase (*ftl*), methenyl-THF cyclohydrolase (*fch*), and the NADPH-dependent methylene-THF dehydrogenase (*mtdA*), all from *M. extorquens* ^53^ (**Figure 6A**, **left**). We previously saw that *fdh*-mediated regeneration of NADH was crucial for malate accumulation from the rTCA pathway. Accordingly, we prototyped a polyphosphate kinase (*ppk*) ATP regeneration system and a NADPH regenerating system employing *fdh* mutants with engineered specificity towards NADPH to regenerate cofactors consumed by the formate assimilation pathway. To generate the biocompatible C1 donor 5,10-CH_2_-THF *in situ*, we coupled the reductive TCA module with the three-enzyme formate assimilation pathway. In total, this represents an eight-enzyme pathway for malate production, all directly expressed from DNA in a one-pot reaction. Previously, we have assimilated formate into glycine and serine using the formate fixing module from *M. extorquens* in combination with the reverse glycine cleavage complex ^29^. In this study, overexpression of the *M.extorquens* genes *ftl*, *fch* and *mtdA* for the formate assimilation pathway in the CFE improved 5,10-CH_2_-THF accumulation 1.6-fold compared to lysate alone, reaching 178 ± 35 μM from 1 mM THF and 10 mM formate after four hours (**Figure 6A**).

**Figure 6.**
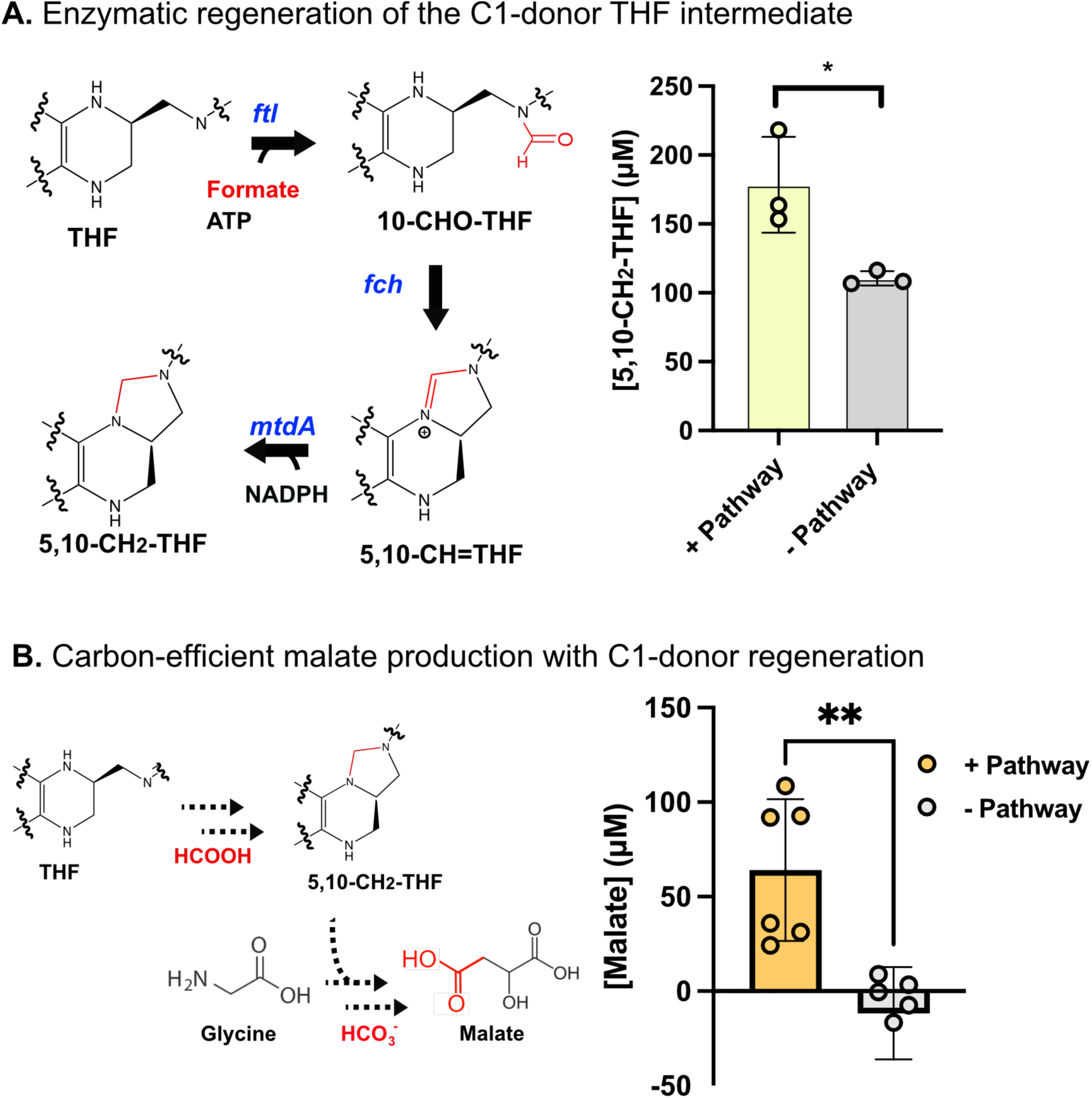
Regeneration of THF cofactor drives carbon-efficient malate production. A. Enzymatic pathway for formate assimilation into THF to form 5,10-CH_2_-THF. The pathway consists of three enzymes and utilizes one ATP and one NADPH equivalent. THF can be recycled back into 5,10-CH_2_-THF after C1 transfer to glycine. 5,10-CH_2_-THF concentration is shown for reactions fed with THF, formate, NADPH, and ATP. Reactions contain CFE either with or without the formate assimilation pathway. Values represent the mean ± standard deviation of three technical replicates. B. Carbon-efficient cell-free conversion of glycine and formate to malate. The pathway consists of eight enzymes for THF recycling, NADH regeneration, and malate synthesis. Two carbon equivalents are incorporated into malate. Malate concentration is shown for reactions fed with all necessary chemical cofactors and starting from formate and glycine. Reactions contain CFE either with or without the full pathway. In this experiment, cell-free bioproduction reactions were run at 250 μl volume. Values represent the mean ± standard deviation of six technical replicates. For all panels, statistical significance was calculated using two-tailed unpaired Welch’s *t*-tests. Asterisks indicate a statistically significant difference (∗: p-value < 0.05, ∗∗: p-value < 0.005).

To regenerate ATP, we first tested a polyphosphate kinase (*ppk*) from a previously published *Erysipelotrichaceae* bacterium ^55^ using inexpensive polyphosphate as a phosphate donor. We found that, when expressing *ppk* in CFE, we could generate 742 ± 3 μM ATP from 1000 µM AMP and 10 mM hexaphosphate in a four hour reaction **(Figure S10)**. However, when we incorporated *ppk* along with the biosynthetic pathway for serine to malate conversion, we found that ATP regeneration did not improve titers over an eight hour reaction. Therefore, we concluded that ATP availability is not limiting flux through the rTCA pathway in these conditions **(Figure S10).** We tested ATP regeneration in the ATP-dependent formate assimilation pathway under standard conditions (1 mM ATP, 1 mM NADPH, 1 mM THF) and found that 5,10-CH_2_-THF accumulation was abolished, despite the ATP regeneration system itself remaining functional **(Figure S10).** This result indicates that the ATP regeneration system introduces pathway incompatibilities in our system, and so it was excluded in future experiments.

To regenerate NADPH consumed by *mtdA* during formate assimilation, we tested two NADPH-selective *fdh* variants for orthogonal NADH and NADPH regeneration *in situ*. Disappointingly, we were not able to see NADPH accumulation in reactions fed with 1 mM NADP^+^ after screening variants from *A. thaliana* and *Candida methylica* **(Figure S11)** ^56,57^. This may be due to poor expression or low activity of these enzymes, resulting in them not being able to compensate for native NADPH consumption in the lysate. Thus, our final reaction set-up for cell-free malate production consisted of the eight-enzyme system integrating rTCA, formate assimilation, and NADH regeneration (**Figures 1A and 6B**).

We applied the formate assimilation pathway for 5,10-CH_2_-THF regeneration to produce malate from C2 glycine and the C1 compounds formate and bicarbonate directly. We co-expressed the formate assimilation module with the rTCA cycle and *fdh* for NADH regeneration. Reactions fed with 10 mM formate, 10 mM bicarbonate, 1 mM THF, and 1 mM glycine produced 64 ± 37 μM of malate in an eight-hour reaction (**Figure 6B**), which is about half of the malate produced from 1 mM 5,10-CH_2_-THF fed in directly with 1 mM glycine (117 ± 6 μM) (**Figure 5B**). With 2 CO_2_ equivalents fixed per malate produced, integrating the formate assimilation pathway with the rTCA cycle process enabled incorporation of 0.13 moles of CO_2_ equivalents per mole of glycine fed.

### Techno-economic analysis of cell-free bioproduction of malate

Despite a number of recent successes of large-scale cell-free biomanufacturing ^58–60^, there remains a perception that the commercial realization of CFE as a metabolic engineering platform is necessarily limited by the costs of catalyst preparation and chemical requirements. To evaluate the economic feasibility of industrializing cell-free bioproduction, we used our platform for malate bioproduction from formate, bicarbonate, and glycine as a basis for performing techno-economic analysis (TEA). TEA is a valuable tool to provide insights into the challenges and opportunities to reduce costs associated with bioproduction.

The use of low-cost feedstocks and cofactor regeneration systems will be crucial to achieving economically competitive cell-free metabolic engineering ^11^. In our previous work, we have shown that glycine and serine can be produced using cell-free based biocatalysts solely from formate through the rGCV pathway with a yield on formate of up to 30% ^29^. In this work, we showed malate production from serine with a yield of 27% (**Figure 5B**). With further optimization, it should be possible to integrate the rGCV and rTCA pathways in CFE and enable malate production from formate directly. Therefore, our TEA assumes that malate can be produced using formate and bicarbonate as the primary feedstocks, with 20% conversion efficiency of formate as the base assumption. Additionally, we demonstrate that enzymatic recycling of NADH and THF can be used to reduce the need for exogenous cofactors; and thus our TEA assumes a continuous process with the ability to regenerate crucial cofactors *in situ*.

The chemical process block diagram is given in **Figure 7A** and base case assumptions are defined in **Table S2**. Our TEA uses the “n^th^” plant assumption, which assumes that similar plants have already been established and therefore does not account for excess costs associated with a first-time process (**Methods S2)**. The TEA revealed that malate produced from formate via a cell-free process could reach a minimum selling price (MSP) of $9.6/kg with the given assumptions (**Figure 7B**, **Table S3, Methods S2)**. This price is only 5.4-fold higher than the current market price for fossil-fuel derived malate ($1.8/kg) ^62^, suggesting that further technological improvements could result in cost-competitive bioproduction. Additionally, we considered the case where formate and glycine are co-utilized as inputs for the cell-free production of malate. In this case, our analysis showed that the cost of malate production only increased by 8%, up to $10.4/kg (**Figure 7B**).

**Figure 7:**
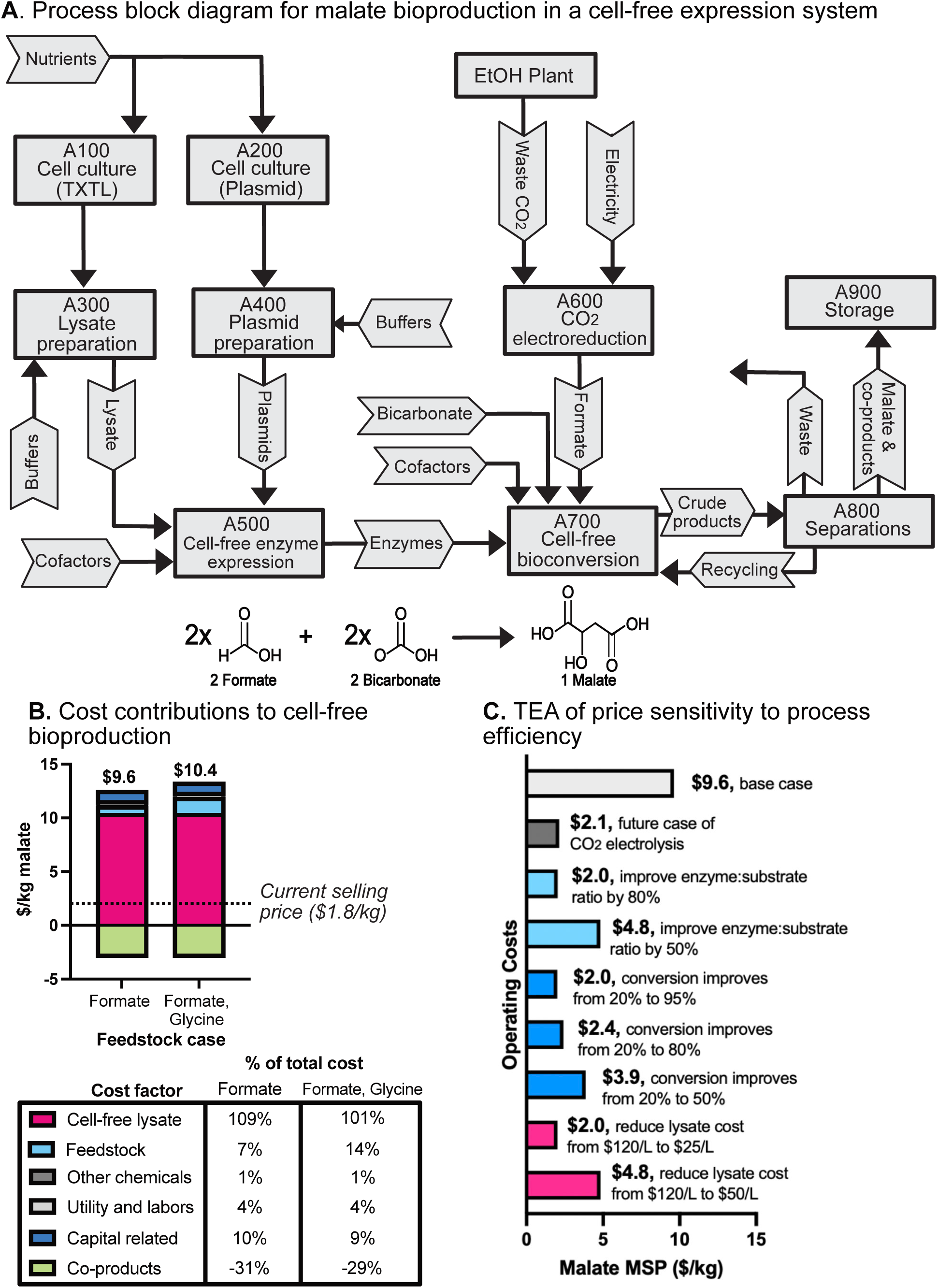
Techno-economic analysis of cell-free bioproduction of malate. A. Process block flow diagram to produce malate by cell free bioconversion. The cell-free bioproduction plant is assumed to be located next to a bio-ethanol plant. Carbon dioxide effluent from the ethanol plant is fed to a low temperature electrolysis process converting the CO_2_ stream to formic acid. An average CO_2_ flow rate of 227 thousand metric ton per year is used to estimate the process scale^88^. Economic analysis and technical assumptions of the CO_2_ conversion step have been established by a multi-national lab team ^64^. Capital and operating costs of the cell free process are assumed to be comparable to the enzyme production process published by NREL ^89^. Cost assumptions of the biological conversion of formic acid to malic acid is taken from previous work and outlined in **Methods S2** ^64^. B. Contributions of different inputs to the cost of cell-free malate bioproduction are shown. Two cases are considered, one where formate is used as the sole carbon source, and one where formate and glycine are co-fed into the bioproduction platform. The co-purification of co-products succinate and citrate are included to offset costs associated with the process. The current selling price of malate produced from fossil fuels ($1.8/kg) is shown with a dashed line. C. Techno-economic analysis of price sensitivity to operating costs and process efficiency. Operating costs, including cost of formate from CO_2_ electrolysis, enzyme turnover, conversion efficiency, and cell-free reagent cost are varied to understand their impacts on the final product cost. TEA suggests that operating costs were the primary drivers affecting malate MSP. The effect of improvements in various operating costs on the final MSP of malate are shown. Assumptions for TEA are given in **Table S2** and **Methods S2**.

We found that operating costs, specifically the cell-free reagent price, are the most significant cost drivers (**Figure 7B**). Process costs of bioproduction can help be offset by co-purifying and selling valuable co-products. In our system, we saw significant accumulation of succinate and citrate (**Figure 3B**), organic acids with large defined markets. Therefore, our analysis calculates the MSP of malate by summing the input costs and subtracting the price of the sellable co-products (**Figure 7B**). The cost of the cell-free lysate at $120/L^11^ accounts for nearly 100% of the process costs, with feedstock accounting for roughly 10% of the total cost (**Figure 7B**). This is in stark contrast to traditional cell-based bioproduction platforms, in which feedstock costs account for up to 60% of total costs ^63^.

Cell-free systems have the potential for near-stoichiometric conversions due to the high control over reaction conditions. Therefore, improving conversion efficiency is a promising strategy to help CFE reach economic viability in coming years. Our experimental results indicated that there is substantial diversion of pyruvate to other metabolites (**Figure 2**), suggesting that conversion efficiencies and the MSP could be improved through lysate engineering and reaction optimization to reduce loss of carbon through competing pathways. Our analysis shows that improving conversion efficiency of this proof-of-concept system to 95% would be needed to enable a competitive MSP (**Figure 7C**).

Beyond optimizing cell-free reaction conditions, improvements in other technical areas will be crucial to lower costs associated with cell-free metabolic engineering and bioproduction from renewable formate. Cell-free lysate preparation is a major driver of the cost for cell-free bioproduction. We found that reducing the cost of lysate preparation from $120/L to $25/L alone could reduce the MSP of malate to $2.0/kg (**Figure 7C**), making it competitive with petroleum-derived production. As the price of labor is the primary cost-driver of cell-free preparation, scaling its production could help reduce associated costs ^12^. The efficiency of electrochemical CO_2_ conversion to formate is expected to increase by up to 50% in coming years ^64,65^, which could further reduce the production cost of malate to $2.1/kg (**Figure 7C**). Metabolic engineering efforts to produce higher value compounds, such as fatty acids derived from acetyl-CoA or other organic acid derivatives, would also permit a higher MSP. While not economically competitive in its current state, this TEA shows promise for the sustainable production of platform chemicals using CFE.

## Discussion

The use of formate and other renewable C1 feedstocks has great potential for sustainable and low-cost bioproduction of commodity chemicals ^1,7,66^. Here, we demonstrate production of the central TCA metabolite, malate, using an “off-the-shelf” CFE without additional genetic engineering. These are widely accessible through commercial vendors and have been broadly adopted for genetic circuit engineering and enzyme overexpression and prototyping ^67–70^. While there are many examples of malate production in microbial systems via the rTCA cycle, these typically still utilize refined glucose as a feedstock and introduce significant growth defects to the engineered strain ^26,71,72^. Additionally, naturally C1-assimilating organisms tend to be difficult to engineer and are limited in their uptake and incorporation of C1 feedstocks ^7^. In this work, we use unique aspects of CFE to enable carbon-conserving production of the industrial di-acid malate that would be very difficult to achieve in microbial cell-based fermentation. Specifically, we were able to 1) minimize background reactions by diluting the CFE, 2) minimize flux through carbon-inefficient oxidative TCA with no strain engineering by introducing chemical inhibitors, and 3) overcome cellular transport limitations to utilize formate as a substrate by using CFE. Collectively, these aspects of CFE allowed us to redirect flux, minimize background reactions, and maintain stable titers of malate while conserving 43% of the carbon that would have been lost otherwise. The approaches developed in this work should be widely applicable to metabolic engineering efforts for other small molecules in CFE. This work establishes a benchmark for the potential of CFE for carbon-conserving prototyping and metabolic engineering.

Decoupled enzyme expression and bioconversion is not a new idea to the field of bioproduction ^73^. For cell-based fermentation, this often involves growing the cells to a sufficient biomass before inducing heterologous protein expression and bioconversion. Traditionally, purified enzyme systems have decoupled these stages by separate expression and enrichment of individual enzymes before combining biocatalysts and chemicals for bioconversion ^14,15,50^. By contrast, we achieve decoupling in a one-pot reaction mixture, in which all enzymes are expressed from DNA in a single reaction before being combined with substrates and cofactors. Additionally, our lysate-based cell-free reaction operates efficiently with roughly 100-fold less enzyme than comparable *in vitro* systems ^14–16^ (**Table 1**).

While one-pot enzyme expression and bioconversion reduces process complexities, it faces the challenge of competition from endogenous enzymes left in the CFE. Enzymes from the TCA cycle have been shown to be highly active in lysates prepared from *E. coli* ^74,75^, creating a challenge for efficiently routing carbon flux into desired products. By utilizing the reductive TCA cycle, one CO_2_ equivalent is incorporated per malate produced from pyruvate. By avoiding decarboxylations through the oxidative TCA cycle, an additional 2 mol CO_2_/mol malate is conserved. Starting from pyruvate, our system reduced carbon loss by an estimated 43% compared to malate that was produced by the endogenous metabolism of the lysate (**Table 1**, **Figure 5B**). We also show that an additional CO_2_ equivalent can be incorporated into malate via formate and glycine by integrating the formate assimilation pathway with the reductive TCA cycle.

Crucial to these accomplishments were the findings that CFE can be diluted following protein expression and that inexpensive inhibitors added directly to reactions to reduce competition from residual enzymes present in the lysate. Specifically, we found that the inhibition of citrate synthase following CFE dilution improved our malate titers 14-fold in a four hour reaction **(Figure S5)**. Dilution of the CFE presents an efficient method to reduce the effect of endogenous enzymes and study engineered pathways that would not be possible in a microbial system. Direct addition of inhibitors into a bioproduction reaction would also likely not be feasible in a microbial host, as the inhibitor may introduce toxicity or be limited in its uptake through the cell membrane. Together, these interventions improved pathway-dependent bioproduction and reduced loss of carbon through CO_2_.

The utility of CFE for rapid prototyping and optimization of *in vivo* metabolic engineering programs has been demonstrated widely ^67,68,76^. The development of CFE as a metabolic engineering platform for carbon-conserving biosynthesis, however, faces many systems-level challenges. For one, the utility of cell-free reactions has been limited by the inability to regenerate necessary cofactors and the loss of activity from pathway enzymes. In our system, we found that malate accumulation could be extended up to 24 hours, compared to 30 minutes, by diluting the CFE following gene expression and regenerating NADH to drive flux towards malate (**Figure 2A**). However, we found that malate titers peaked at 10 hours when starting from serine, and then decreased until reaching baseline levels **(Figure S9),** suggesting that the engineered pathway had lost function, or that the produced malate is siphoned off to other reactions in the cell lysate. We also found that *in situ* ATP regeneration did not improve malate yields in our system **(Figure S10)**, although efficient regeneration of ATP may be necessary to drive flux in other engineered pathways. Broadly, ATP regeneration systems are well studied for cell-free gene expression ^77–79^, while the development of generalizable, cost-effective regeneration systems applicable to biosynthetic pathways are still needed ^11,80^. Traditionally, *in vitro* reactions have relied on supplementing purified enzymes and cofactors throughout the reaction to extend lifetimes ^14,15^. Notably, our system reaches comparable reaction lifetimes to previous *in vitro* systems without the supplementation of additional enzymes and cofactors. Generalizable strategies to improve enzyme stability and regenerate cofactors *in situ* will greatly reduce the barriers to achieving cell-free bioproduction at scale.

Other considerations to optimize engineered pathways in CFE are the balancing of cofactor compatibilities, reaction conditions, and enzyme activities, among other parameters ^8,18,81^. As the number of steps in an engineered pathway increases, incompatibilities with respect to cofactor and reaction condition requirements can often arise ^15,17,43^. In this work, we found that rational screening of enzyme isoforms could help alleviate pathway incompatibilities to construct larger pathways **(Figure 4D, 4E).** Additionally, some degree of reaction screening in different buffers and DNA template ratios was necessary to construct our eight-enzyme pathway **(Figures S12-S14)**. The performance of this system with minimal optimization demonstrates the high potential for carbon-conserving pathway engineering with high efficiency in CFE. A much more exhaustive and systematic survey of reaction conditions would be necessary to truly optimize the pathway performance. While the high dimensionality of the design space makes predicting optimal conditions *a priori* difficult, model-driven engineering may help bridge experimentation and machine learning to accelerate systems-level optimization of bioproduction platforms ^82–84^.

For our base case TEA analysis, we assumed 100% cofactor recycling, which otherwise accounts for 97% of the cost for a bench-top scale CFE conversion of formate to serine ^29^. Significant improvements in cofactor regeneration will be needed to be able to negate such large costs. The drop in malate concentrations after 10 hours **(Figure S9)** signifies the need for additional product removal unit operations for a successful transition from batch production to a continuous production modelled in our TEA analysis. Taking these caveats into account, our work demonstrates an innovative carbon-conserving alternative for large scale bioproduction of malic acid.

The global malic acid market is estimated to be around 100,000 metric tons annually, produced almost entirely from petrochemicals ^85^. Cell-free production of malate using electrochemically generated formate has the potential to reach a higher carbon efficiency than microbial production, which has failed to achieve commercial scale to date. Less than 10% of all CO_2_ waste streams generated by U.S. bioethanol plants would be needed to satisfy the global malic acid market, assuming just a 5% yield on formate ^86^. In turn, this would avoid roughly 400,000 tons of CO_2_ emissions annually associated with traditional synthesis ^87^. Our work demonstrates construction of carbon negative pathways by assimilation of formate and bicarbonate, as well as carbon conservation steps in the rTCA cycle to produce central metabolites such as pyruvate and malate, which themselves can be a starting point for other metabolic pathways. Cell free biocatalysis allows metabolic pathways to be stitched together in a modular nature by addition of appropriate enzymes. Pyruvate, an intermediate in our engineered pathway, can be routed through acetyl-coA to produce a variety of commodity chemicals such as fatty acids, terpenes, alcohols or polymers ^19, 80, 84^. Given the rapid growth of the field in recent years, cell-free bioproduction from renewable formate may soon be able to address key limitations of microbial fermentation and replace petroleum-based chemical production.

## Methods

### 1. Plasmid Preparation

Plasmids expressing pathway genes were cloned in *E. coli* NEB Turbo cells. All PCR amplification of genomic DNA used Phusion DNA polymerase. Primers were synthesized by IDT and gBlocks were synthesized by Twist Biosciences. Both primers and gBlocks were resuspended with nuclease-free water. Plasmid assembly was achieved using 5X In-Fusion HD mastermix (Takara). Assembled plasmids were plated onto LB-agar plates with 100 μg/mL carbenicillin. Transformed cells were grown overnight at 30 °C. Single colonies were picked from plates and grown overnight in LB shaking at 30 °C with 100 μg/mL carbenicillin.

Plasmids were isolated from subcultures using a DNA miniprep kit (QIAprep Spin Miniprep Kit) and sequenced with Sanger (Genewiz inc.) or full-plasmid sequencing (Primordium) to verify correctly assembled plasmids. Plasmids were grown in culture volumes of ∼100 mL to ensure adequate yields for multiple cell-free reactions. Plasmids were further purified using a PCR purification kit (Invitrogen PureLink, Cat. K310001) and eluted with nuclease-free water. Plasmid concentrations were quantified via spectrophotometry (Nanodrop 2000c, Cat. ND-2000C).

### 2. Cell-Free Protein Synthesis Reactions

The cell-free system was acquired from Arbor Biosciences (myTXTL). The cell-free system used for an experiment was thawed on ice and pooled into a 1.5 ml Eppendorf tube, vortexed, and spun-down using a mini benchtop centrifuge to ensure homogeneity across samples. For reactions containing three or fewer genes, reactions were assembled on ice from the CFE, purified DNA, and necessary cofactors. The CFE was pipette mixed and added to each PCR tube in 7.5 μL for a final volume of 10 μL. These PCR tubes were incubated overnight at 30C. For reactions involving more than three genes, plasmids and cofactors were mixed with an acoustic liquid handler robot (Echo Labcyte 525) into Labcyte 384-well destination plates (001-14555). The 384-well plates were then incubated at 30°C overnight.

### 3. Cell-Free Bioproduction Reactions

Cell-free bioproduction reactions were mixed in 25 μL containing 2.5 μL of CFE-expressed enzymes. CFE-expressed enzymes were diluted in 10 mM Tris pH 8 prior to adding if they were diluted beyond 10-fold in the final reaction. Bioproduction reactions were done in 50 mM HEPES pH 8. For reactions containing three or fewer enzymes, reactions were assembled by hand from the CFE-expressed enzymes and necessary substrates and cofactors. For reactions involving more than three enzymes, enzymes and chemicals were mixed with an acoustic liquid handler robot (Echo Labcyte 525) into 96-well V-bottom plates (Costar, Cat. 3363). The plates were sealed with a foil adhesive (Thermo, AB0626) and the reactions were run for 4 -8 hours at room temperature.

### 4. Metabolite quantification with LC/MS

Samples were analyzed via Agilent 6530 LC/Q-TOF in negative mode using a BEH Amide 50 mm column (Waters, 186004800). Standard curves were prepared by spiking known amounts of metabolites into diluted CFE and HEPES. For LC/MS, the aqueous phase was LC/MS grade water and the organic phase was 95/5 acetonitrile/water with 10mM ammonium acetate and .04% v/v ammonium hydroxide. The % aqueous/organic gradient was run as follows: hold at 5/95 for 2.5 minutes, move to 33.5/66.5 over 5 minutes, 40/60 over 1 minute, hold 40/60 for 1 minute, then return to 5/95 over 1 minute. The flow rate was held at 0.5 mL/min.

### 5. Statistics

Statistical significance was calculated using two-tailed unpaired Welch’s *t*-tests. Asterisks in Figures indicate a statistically significant difference (∗: p-value < 0.05, ∗∗: p-value < 0.005).

