## Supplementary Information for "Carbon-conserving Bioproduction of Malate in an *E. coli*-based Cell-Free System"

**Supplementary Figures**


**Figure S1:** Thermodynamics for the synthesis of malate from glycine

**Figure S2:** *mdh*-dependent malate concentrations over time

**Figure S3:** Proteome analysis of TCA enzymes in an *E. coli*-based CFE

**Figure S4:** Effect of dilution on *mdh*-dependent malate production

**Figure S5:** Time-course measurements of interventions to improve *pyc*-dependent malate production

**Figure S6:** Hydroxycitrate-dependent acetyl-CoA accumulation

**Figure S7:** Validation of glyoxylate shunt as a complementary carbon-conserving pathway in CFE

**Figure S8:** Dilution of CFE improves pathway-specific pyruvate production

**Figure S9:** Time-course serine to malate bioconversion via a four-enzyme pathway

**Figure S10:** Effect of ATP-regenerating ppk on cell-free biosynthesis

**Figure S11:** Lack of activity from NADPH-specific *fdh* variants in CFE

**Figure S12:** Pyruvate accumulation in the presence of downstream cofactors

**Figure S13:** Effect of competing DNA expression on *tdcB* activity

**Figure S14:** Effect of titrating plasmid DNA on bioconversion efficiency

**Supplementary Tables**

**Table S1:** Bioproduction & carbon efficiency calculations

**Table S2:** Base case TEA assumptions

**Table S3:** Base case TEA results

**Table S4:** DNA sequences used in this study

**Supplementary Methods**

**Methods S1:** Comparative sequence analysis of enzyme isoforms

**Methods S2:** Techno-economic Analysis

**Supplementary References**

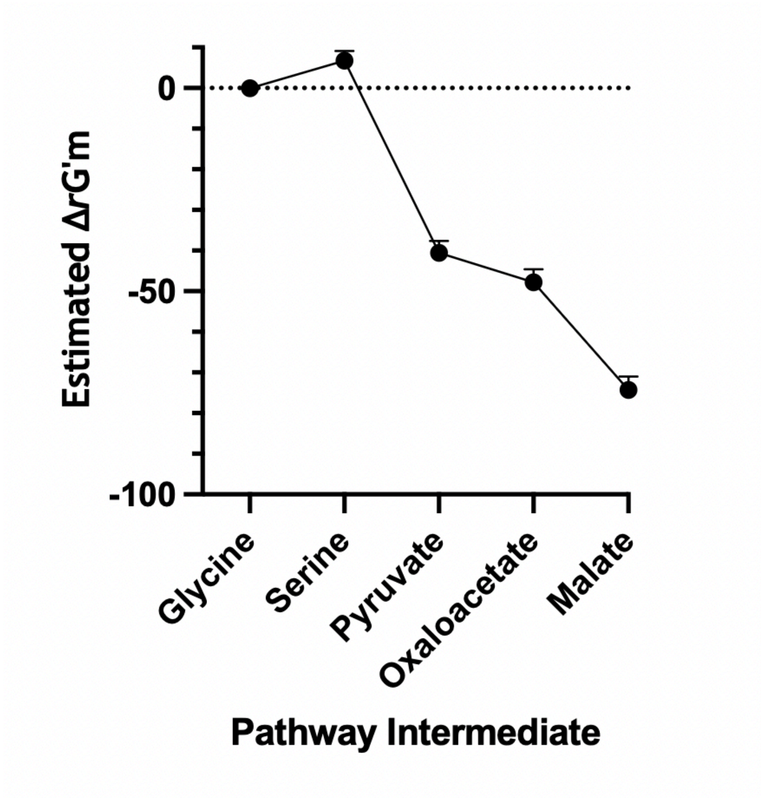


<u>Figure S1. Thermodynamics for the synthesis of malate from glycine.</u> The ΔG'° of each step was calculated using eQuilibriator assuming a standard concentration of 1mM for all reactants.

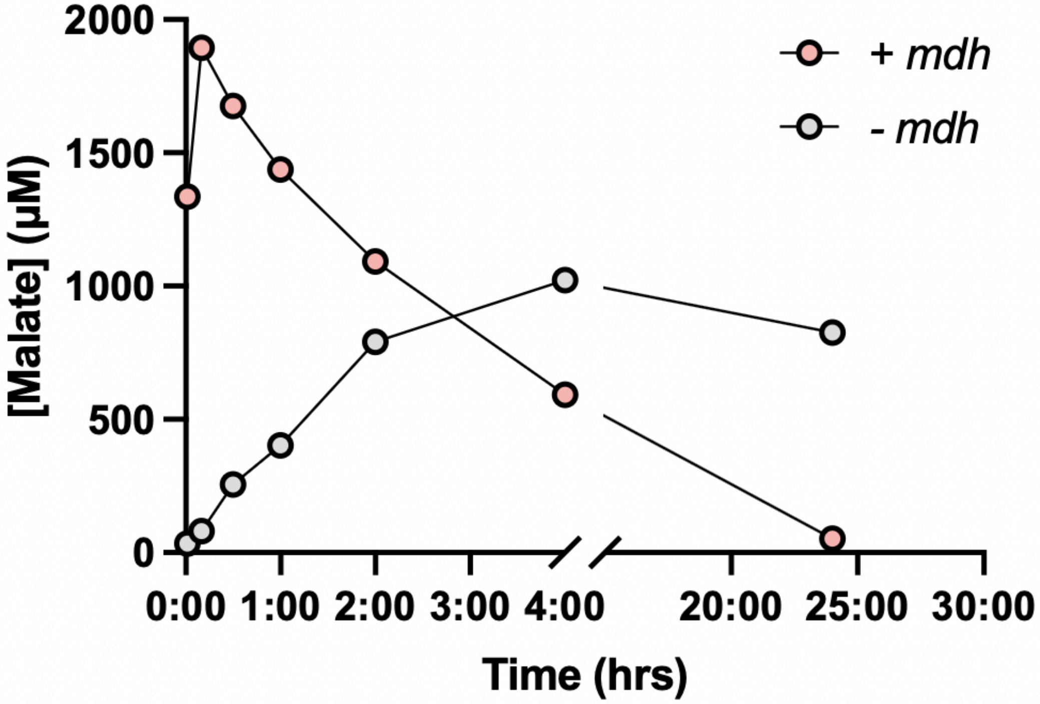


Figure S2. *mdh*-dependent malate concentrations over time. Malate concentrations are shown over 24 hours for CFE with or without *mdh* overexpression. Reactions contain oxaloacetate and NADH at 1mM each. Time points were collected at 0, 10, 30, 60, 120, 240, and 1440 minutes.

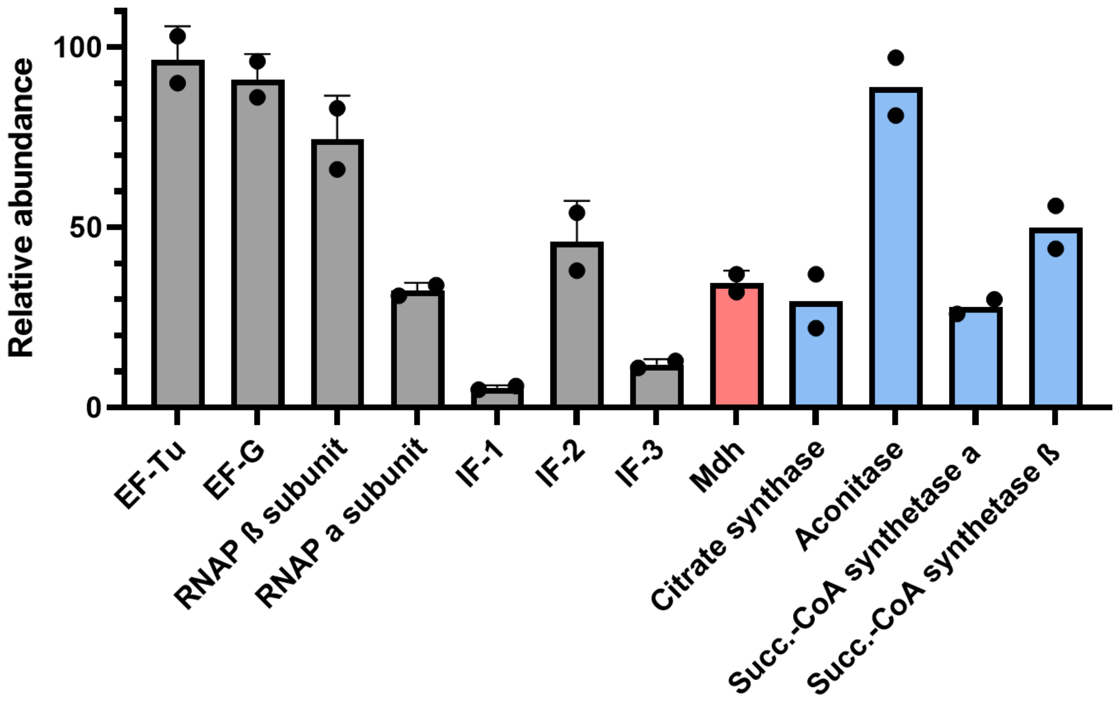


Figure S3. Proteome analysis of TCA enzymes in an *E. coli*-based CFE. Published proteomic data from CFE was analyzed to compare relative amounts of TCA cycle enzymes to highly expressed enzymes required for key transcription and translation processes.

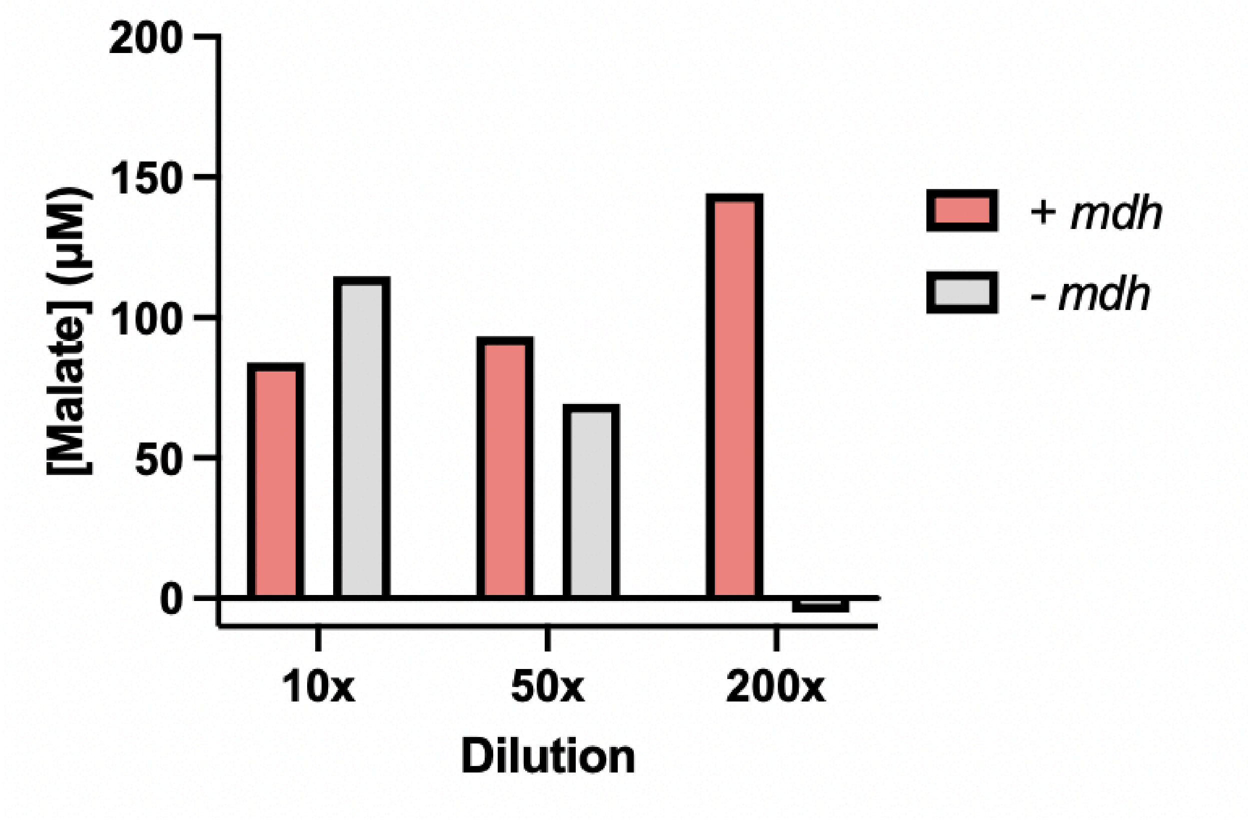


<u>Figure S4. Effect of dilution on *mdh*-dependent malate production.</u> Malate concentrations are shown for reactions expressing *mdh* and *fdh* or *fdh* alone. Reactions were diluted 10, 50, or 200-fold into a buffer containing 1mM NADH, 10mM formate, and 1mM oxaloacetate. Malate concentrations were measured after four hours.

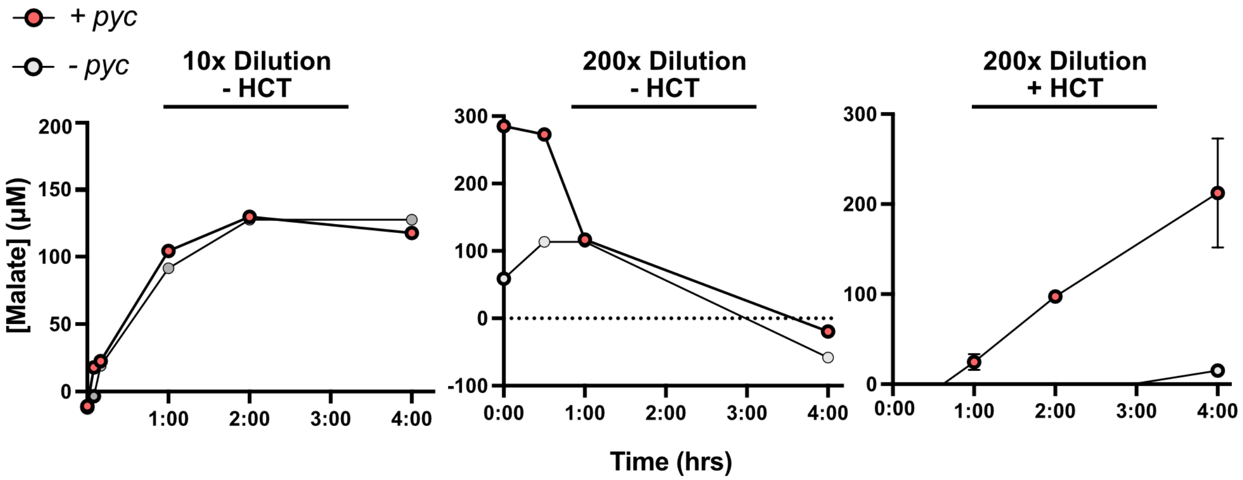


Figure S5. Time-course measurements of interventions to improve *pyc*-dependent malate production. Malate concentrations are shown over four hours for CFE reactions diluted either 10- or 200-fold and with or without hydroxycitrate (HCT) as an inhibitor of the TCA cycle. Reactions in the left panel contain 0.5mM pyruvate, 1mM $HCO_3^-$, 1mM ATP, 1mM $Mg(CH_3COO)_2$, 1mM acetyl-CoA, 1mM NADH, 10mM formate. Reactions in middle and right panels contain 1mM pyruvate, 5mM $HCO_3^-$, 1mM ATP, 2mM $Mg(CH_3COO)_2$, 1mM acetyl-CoA, 1mM NADH, 15mM formate. Reactions containing HCT contain 2.5mM.

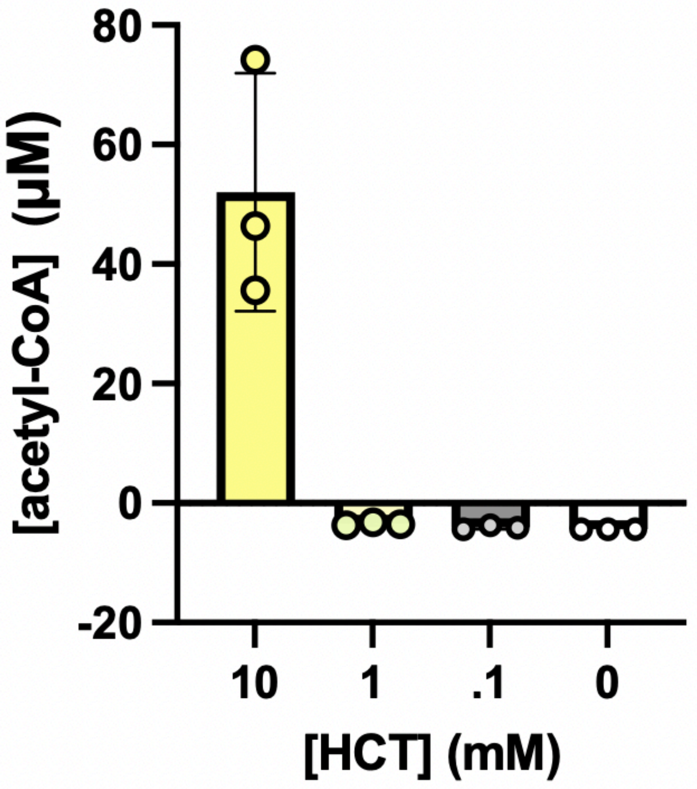


Figure S6. Hydroxycitrate-dependent acetyl-CoA accumulation. Acetyl-CoA concentrations are shown after four hours for reactions with different concentrations of hydroxycitrate (HCT). Reactions express *pyc*, *mdh*, and *fdh*, and contain all of the needed substrates and cofactors for pyruvate to malate conversion. Acetyl-CoA is a substrate for citrate synthase, along with the product of *pyc*, oxaloacetate. The accumulation of acetyl-CoA at 10mM of HCT suggests that citrate synthase is partially inhibited at this concentration. Values represent the mean ± standard deviation of three technical replicates.

A.

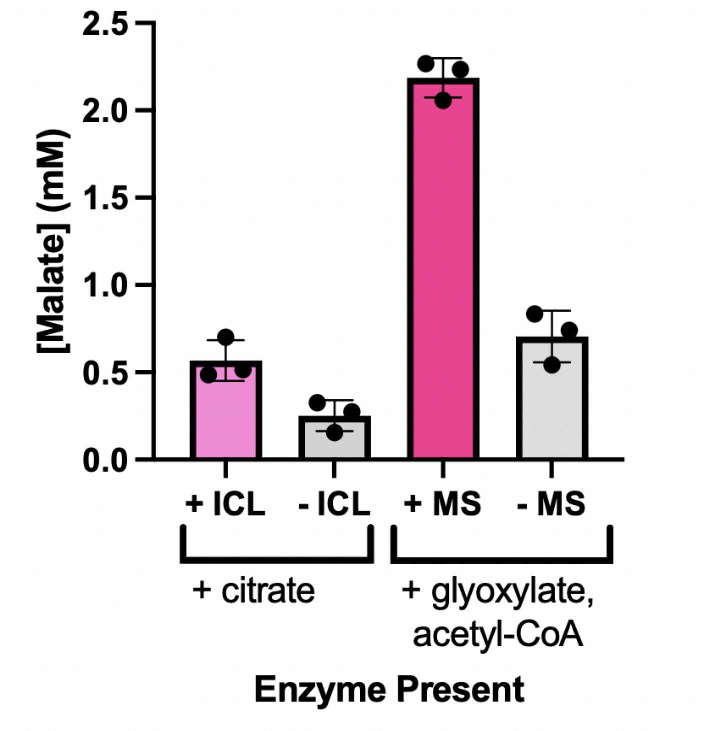
2.5
2.0
1.5
1.0
0.5
0.0
[Malate] (mM)
+ ICL
- ICL
+ MS
- MS
+ citrate
+ glyoxylate,
acetyl-CoA
Enzyme Present

B.

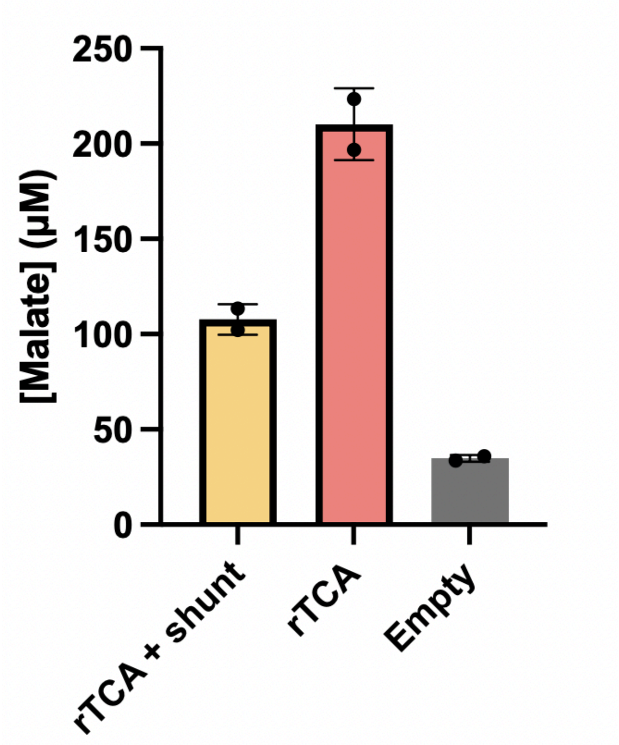
250
200
150
100
50
0
[Malate] (μM)
rTCA + shunt
rTCA
Empty

Figure S7. Validation of glyoxylate shunt as a complementary carbon-conserving pathway in CFE. Concentrations of malate are shown with different variations of the glyoxylate shunt expressed.

**A.** Isocitrate lyase (ICL) and malate synthase (MS) were individually expressed in CFE and mixed in reactions with their respective substrates. Malate concentrations were measured after four hours to verify the individual activity of these enzymes in CFE.

**B.** Reactions containing either the rTCA module (*pyc*, *mdh*, and *fdh*) or rTCA + shunt (*pyc*, *mdh*, *fdh*, ICL, MS) were measured after four hours. Malate concentrations may be lower in the condition with the shunt expressed due to the limited gene expression capacity or added consumption of necessary intermediates from the shunt pathway. For all panels, values represent the mean ± standard deviation of at least two technical replicates.

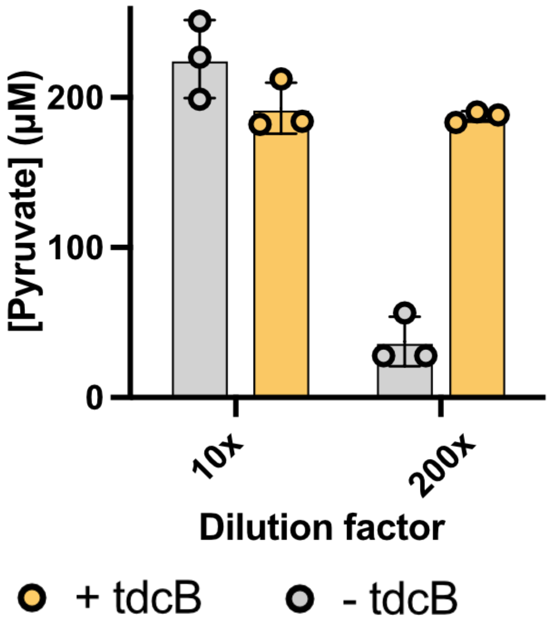


Figure S8: Dilution of CFE improves pathway-specific pyruvate production. Pyruvate concentrations are shown for reactions containing serine, PLP, and AMP. CFE reactions containing *tdcB* or a no enzyme control are diluted 10- or 200-fold in the final reaction. Pyruvate is measured after four hours. Values represent the mean ± standard deviation of three technical replicates.

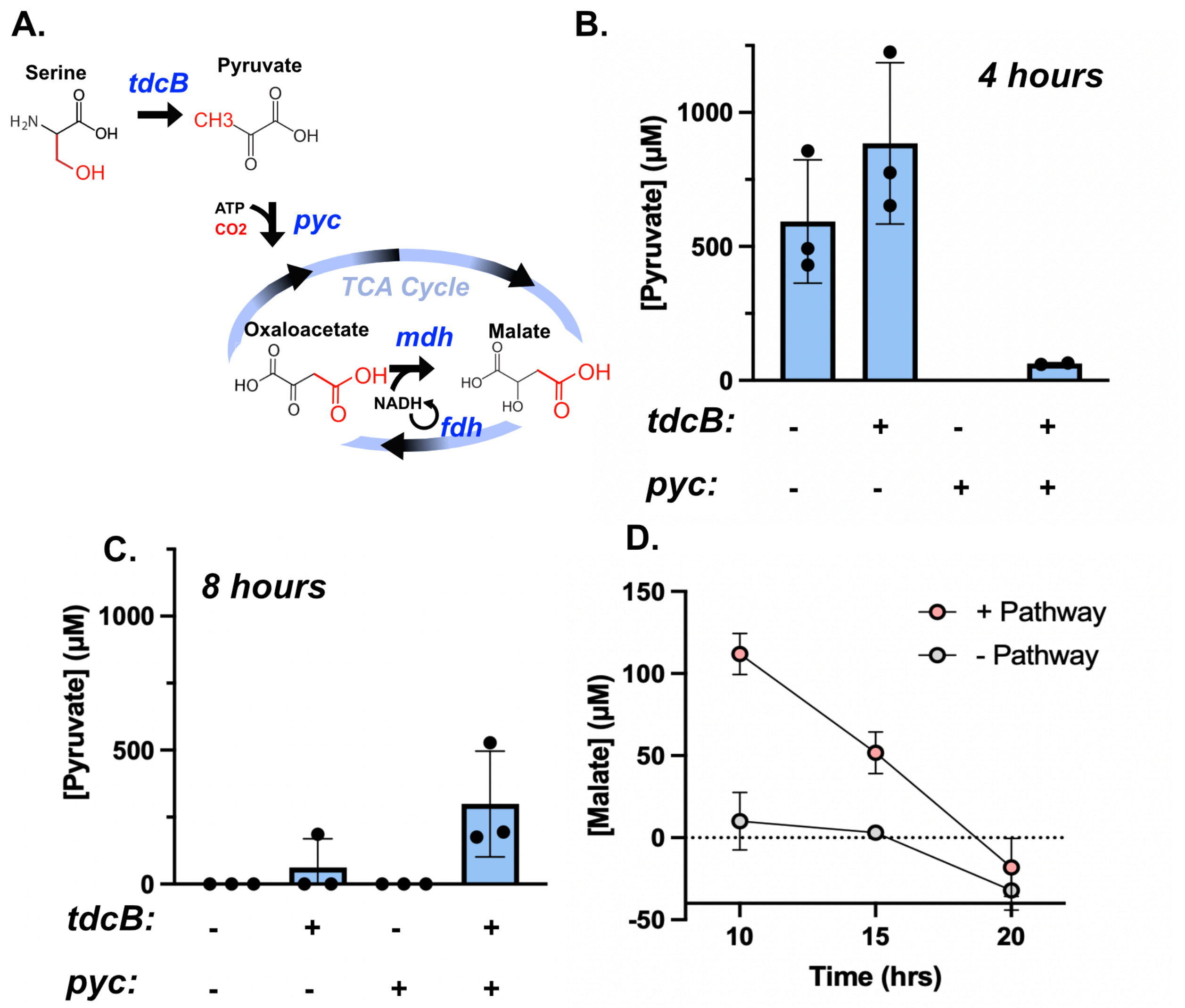


Figure S9: Time-course serine to malate bioconversion via a four-enzyme pathway. **A.** Serine is converted to malate using a four enzyme pathway in which *tdcB* converts serine to pyruvate, and pyruvate is carboxylated into oxaloacetate via *pyc*. Oxaloacetate is then reduced to malate by *mdh*, with cofactor regeneration from *fdh* to drive flux towards malate. For all panels, reactions contain *mdh* and *fdh* expressed at .1 and 1.5 nM, respectively. *pyc* and *tdcB* are either included at 2.5 nM or excluded in each condition. **B.** Pyruvate concentrations are shown at four hours. In reactions with *pyc*, we see a large decrease in pyruvate titers, suggesting it is being converted to oxaloacetate.

**C.** Pyruvate concentrations are shown at eight hours. The buildup of pyruvate in the conditions with *tdcB* expressed suggests that conversion from pyc may not be able to match the rate of production from *tdcB*. **D.** Malate concentrations are shown at 10, 15, and 20 hours when the pathway is expressed. This data is from a different experiment to that shown in panels B and C. The decrease in malate over time may be due to instability of cofactors or enzymes in our engineered pathway. For all panels, values represent the mean ± standard deviation of at least two technical replicates.

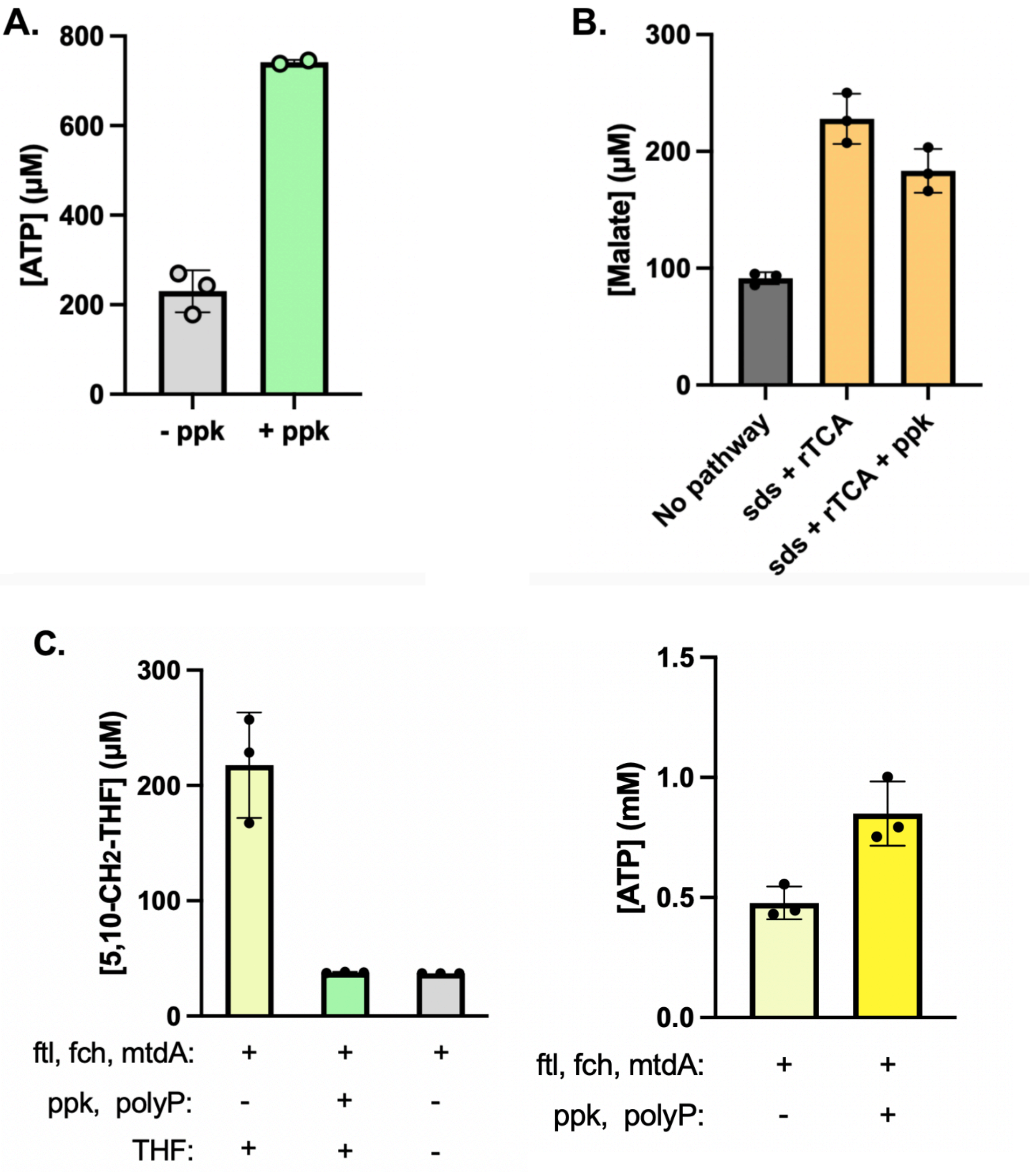
A.
[ATP] (μM)
800
600
400
200
0
- ppk
+ ppk
B.
[Malate] (μM)
300
200
100
0
No pathway
sds + rTCA
sds + rTCA + ppk
C.
[5,10-CH2-THF] (μM)
300
200
100
0
ftl, fch, mtdA: + + +
ppk, polyP: - + -
THF: + + -
[ATP] (mM)
1.5
1.0
0.5
0.0
ftl, fch, mtdA: + +
ppk, polyP: - +

Figure S10. Effect of ATP-regenerating ppk on cell-free biosynthesis. **A.** ATP concentrations are shown for reactions containing polyphosphate kinase from a *Erysipelotrichaceae* bacterium (*ppk12*). Reactions contain 1mM AMP and 10mM polyphosphate. Reactions are collected after four hours. **B.** Malate concentrations are shown for reactions containing sds + rTCA (*pyc*, *mdh*, and *fdh*) with or without ppk. The similar malate concentrations between the conditions with or without ppk suggest that ATP availability may not be limiting in our system for malate production. **C.** 5,10-$CH_2$-THF and ATP concentrations are shown for reactions containing the formate assimilation pathway, consisting of the ftl, fch, and mtdA enzymes, and either including or excluding the ATP regeneration pathway. On the left, reactions either include or exclude the ATP regeneration system shown in Panel A, or THF as a negative control. The loss of 5,10-$CH_2$-THF accumulation in the presence of the ATP regeneration system suggests that it may interfere with function of the formate assimilation pathway. Panels A, B, and C represent separate experiments. For all panels, values represent the mean ± standard deviation of at least two technical replicates.

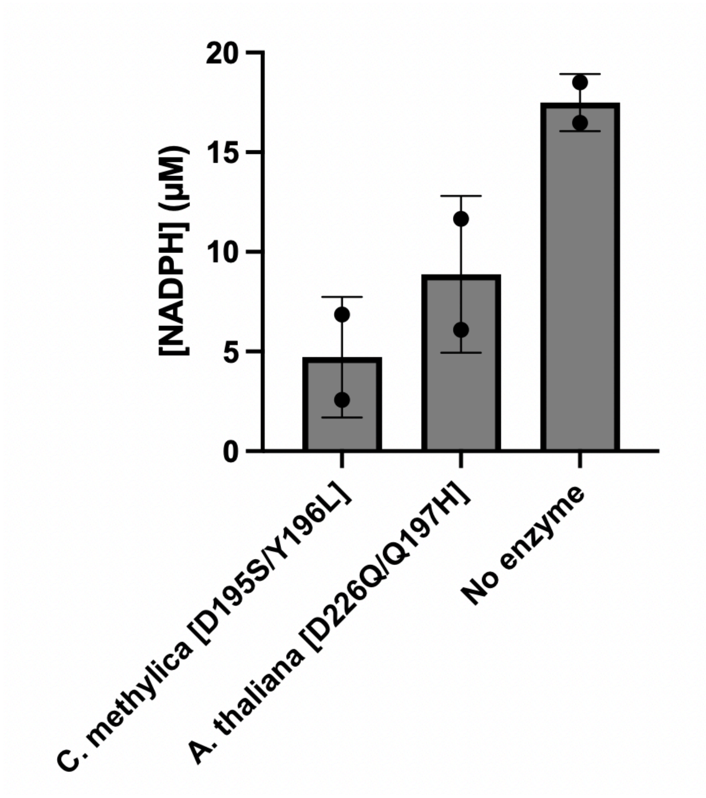


Figure S11. Lack of activity from NADPH-specific *fdh* variants in CFE. NADPH concentrations are shown for three CFE reactions containing either a NADPH-specific formate dehydrogenase (fdh) variant or no enzyme. We did not observe differential NADPH levels for either enzyme tested. This may suggest that these enzymes are not well-expressed in our system, or that NADPH is being consumed rapidly by endogenous enzymes in the CFE. Reactions are fed with 1mM $NADP^+$ and 10mM formate. Reactions are measured after four hours. Values represent the mean ± standard deviation of two technical replicates.

**A.**

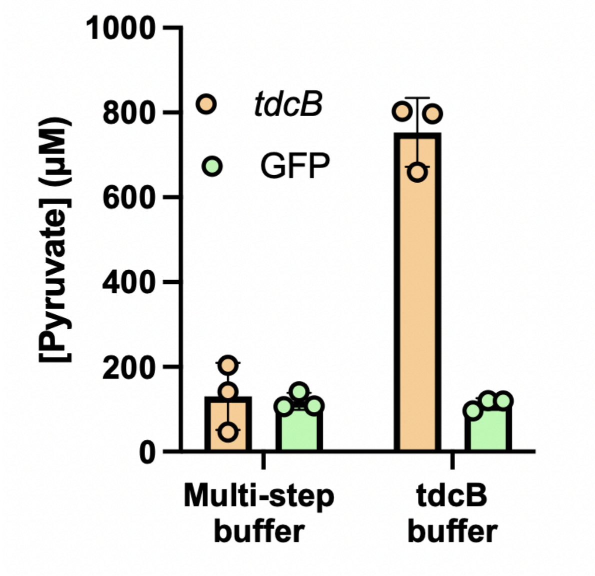


**B.**

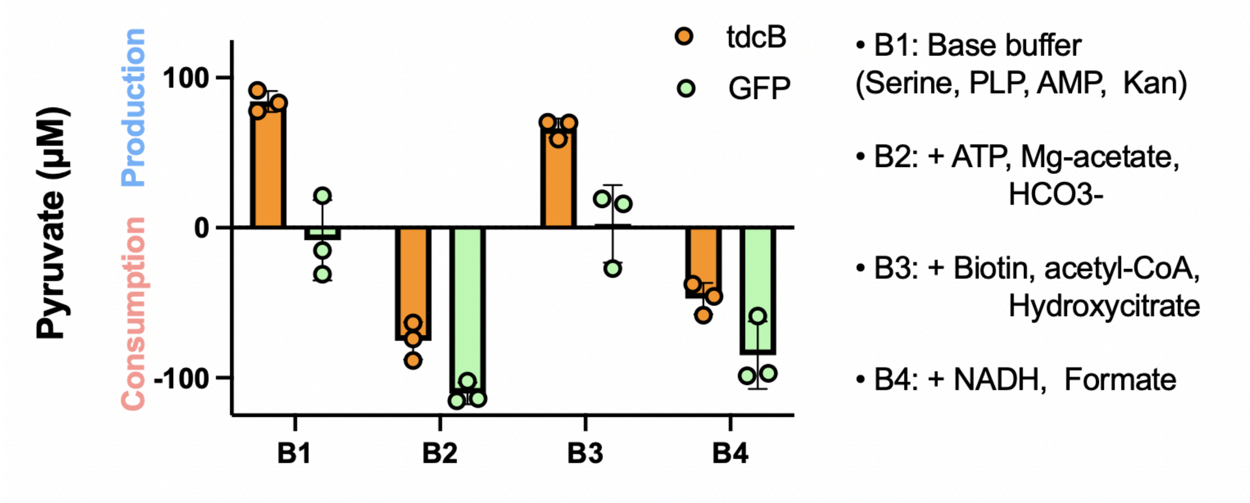


<u>Figure S12: Pyruvate accumulation in the presence of downstream cofactors.</u> To investigate potential pathway incompatibilities, we measured pyruvate accumulation from serine using *tdcB* in two buffer conditions. Pyruvate concentrations are shown in the presence of the cofactors required for downstream steps of the pathway.

**A.** CFE expressing *tdcB* is compared to a reaction expressing GFP in conditions containing either *tdcB* buffer (TB) (1mM AMP, 1mM PLP) or with a multi-step buffer (MSB) with the cofactors required for *pyc* and *mdh* as well (1mM AMP, 1mM PLP, 1mM

ATP, 2mM $Mg(CH_3COO)_2$, 1mM biotin, 1mM acetyl-CoA, 2.5mM hydroxycitrate, 1mM NADH, 10mM formate). Under MSB conditions, we see no significant pyruvate accumulation after four hours.

**B.** Additional buffer conditions are tested to understand which cofactors lead to higher consumption of pyruvate in our system, with each buffer containing different combinations of cofactors required for either *mdh* or *pyc* function. B1 contains the necessary cofactors for *tdcB* function (1mM AMP, 1mM PLP). B2 contains B1 with 1mM ATP, 2mM $Mg(CH_3COO)_2$, and 10mM $HCO_3^-$. B3 contains B1 with 1mM biotin, 1mM acetyl-CoA, and 2.5mM hydroxycitrate. B4 contains B1 with 1mM NADH and 10mM formate. As our CFE contains a significant amount of pyruvate initially, production and consumption are measured relative to the amount of pyruvate present at 0 hours. We found that pyruvate was being consumed faster than it was produced in the buffers containing ATP (B2) and NADH (B4), but not in B1 and B3. These results suggest that the addition of certain compounds that are broadly used throughout metabolism may accelerate the siphoning of pyruvate into side products.

For both panels, reactions are measured after four hours. Values represent the mean ± standard deviation of three technical replicates.

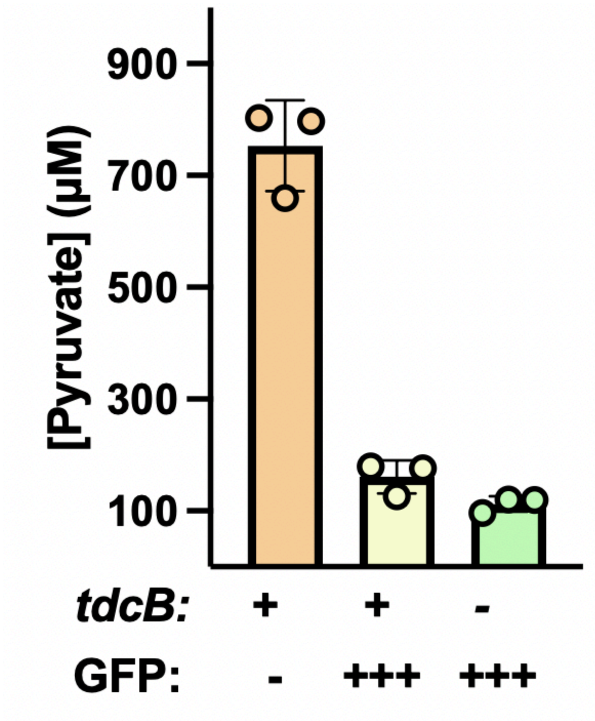


Figure S13: Effect of competing DNA expression on *tdcB* activity. In one-pot cell-free gene expression reactions, there is a tradeoff between the relative amounts of protein that can be expressed from different DNA templates. We demonstrated this tradeoff by measuring pyruvate accumulation via *tdcB* when expressed with or without deGFP as a competing plasmid. Pyruvate concentrations are shown for reactions containing plasmid DNA for *tdcB*, GFP, or both. For reactions with *tdcB*, plasmid DNA is added at 3 nM. GFP plasmid is added at 9 nM. We found that pyruvate production levels from *tdcB* were highly sensitive to the amount of deGFP plasmid added in the system. These results highlight that high expression requirements must be balanced with the total amount of DNA added to the system so that each enzyme can be expressed at relevant levels.

Reactions are measured after four hours. Values represent the mean ± standard deviation of three technical replicates.

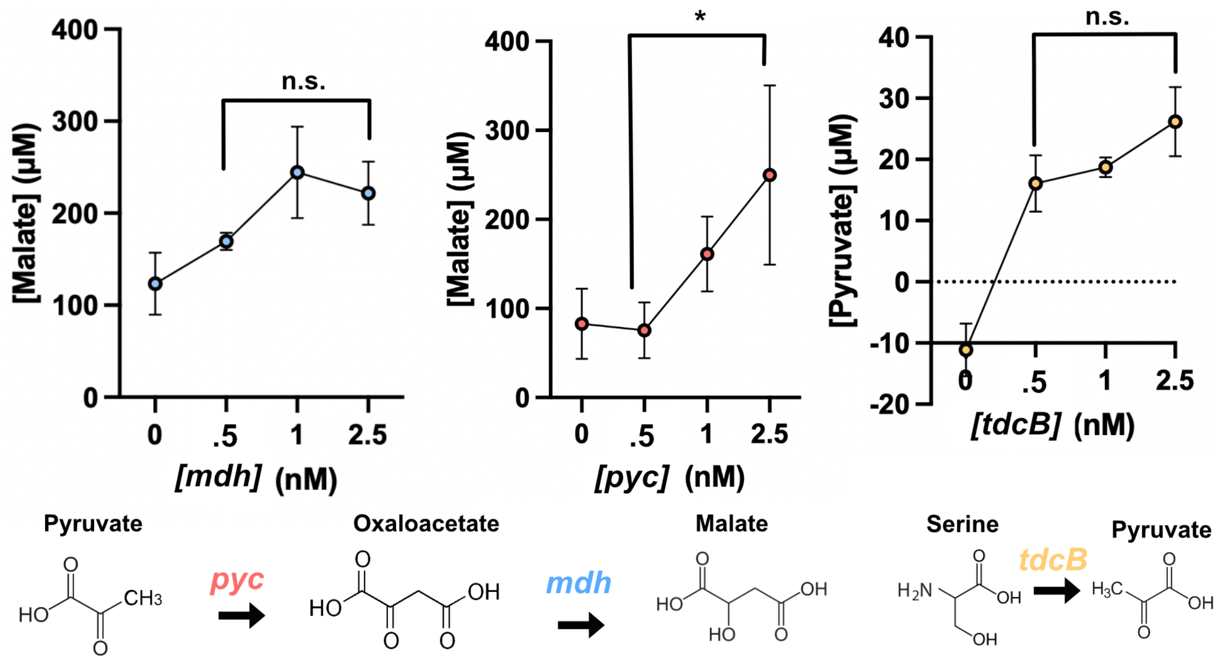


<u>Figure S14: Effect of titrating plasmid DNA on bioconversion efficiency.</u> Product concentrations are shown for reactions titrating either *mdh*, *pyc*, or *tdcB* plasmid DNA to understand which reaction steps were most sensitive to enzyme concentrations. For the *mdh* and *pyc* titrations, *fdh* is held constant at 1 nM. *mdh* was held constant at 1 nM for the *pyc* titration and *pyc* was held constant at 2.5 nM for the *mdh* titration. For both, pyruvate was used as the feedstock at 1 mM with all of the relevant cofactors provided. For the *tdcB* titration, the single step transformation of serine to pyruvate was measured and no other enzymes were expressed. For the single-step *mdh* and *tdcB* reactions, we found that product titers were not significantly different over a five-fold increase in plasmid concentration. These results indicate that low concentrations of enzyme-encoding DNA are sufficient for these reactions, and expression resources can be allocated towards other enzymes. In the *pyc* titration, malate continued to increase

up to 2.5 mM of plasmid DNA, suggesting that *pyc* concentration may be limiting in our system and should be highly expressed.

Reactions are measured after eight hours. Values represent the mean ± standard deviation of at least two technical replicates.

**Supplementary Tables**

Table S1: Bioproduction & carbon efficiency calculations

A. Effect of dilution & inhibition on malate production (Figure 3B)

| | **- pathway<br>- dilution<br>- inhibitor** | **+ pathway<br>- dilution<br>- inhibitor** | **- pathway<br>+ dilution<br>+ inhibitor** | **+ pathway<br>+ dilution<br>+ inhibitor** |
|---|---|---|---|---|
| Malate (μM) | 308 | 348 | 50 | 125 |
| Production from pathway | $\frac{348}{308} = 1.1$ | | $\frac{125}{50} = 2.5$ | |
| Reduction in background reactions | $\frac{308}{50} = 6.2$ | | | |

B. Malate yield on formate (Figure 6B)

| **Malate concentration (mM)** | **Malate concentration (g/L)** | **Formate concentration (mM)** | **Formate concentration (g/L)** | **Yield (g/g)** |
|---|---|---|---|---|
| .064 | .0085 | 10 | .46 | .018 |

C. Malate yield on glycine (Figure 6B)

| **Malate concentration (mM)** | **Malate concentration (g/L)** | **Glycine concentration (mM)** | **Glycine concentration (g/L)** | **Yield (g/g)** |
|---|---|---|---|---|
| .064 | .0085 | 1 | .087 | .098 |

D. Carbon fixation efficiency (Figure 6B)

| Malate through pathway (mM) | Malate without pathway (mM) | Difference in malate production (mM) | CO2/malate (mol/mol) | CO2 fixation (mM) | Maximum CO2 fixation (mM) | $CO_2$ fixation efficiency (%) |
|---|---|---|---|---|---|---|
| .064 | -.011 | .075 | 2 | .150 | 2 | 7.5 |

E. Enzyme usage

| TXTL expression capacity (mg/ml) | TXTL dilution in final reaction vol. | Total expressed enzyme concentration (mg/ml) | Concentration from Luo 2023 (mg/ml) | Concentration from Luo 2022 (mg/ml) | Concentration from Schwander 2016 (mg/ml) |
|---|---|---|---|---|---|
| 4 | 200 | .02 | 10.5 | 10.4 | 3.0 |

Table S2: Base case TEA assumptions [1–3]

| | |
|---|---|
| Formic acid production via CO2 electrolysis | $CO_2$ Electrolysis Current Density, 140 mA/cm<br><br>Onstream Factor, 40%<br><br>Electricity cost, $0.068/kWh<br><br>$CO_2$ Single-Pass Conversion, 20%<br><br>$CO_2$ Electrolysis Cell Voltage, 3.5 V<br><br>$CO_2$ Price, $40/metric ton $CO_2$<br><br>Formic Acid Faradaic Efficiency, 94% |
| Enzyme to formic acid ratio | 1.031 (by mass) |
| Formic acid conversion | 20% |
| Cell free lysate cost | $120/L |
| Enzyme yield | 44.75 g/L |

Table S3: Base case TEA results

| | |
|---|---|
| Total project investment, TPI<br><br>(Excluding the electrolysis process) | MM$ 724 |
| Uninstalled equipment<br><br>(Excluding the electrolysis process) | MM$ 192 |
| Malic acid production | 36,510 metric tons/year |
| Calculated malic acid price using 2020 US$ | $9.63/kg |

Table S4: DNA sequences used in this study

| Gene | Sequence |
|---|---|
| mdh_Ec | ATGAAAGTCGCAGTCCTCGGCGCTGCTGGCGGTATTGGCCAGGCGCTT<br>GCACTACTGTTAAAAACCCAACTGCCTTCAGGTTCAGAACTCTCTCTG<br>TATGATATCGCTCCTGTGACTCCCGGTGTGGCTGTCGATCTGAGCCAT<br>ATCCCTACTGCTGTGAAAATCAAAGGTTTTTCTGGTGAAGATGCGACT<br>CCGGCGCTGGAAGGCGCAGATGTCGTTCTTATCTCTGCAGGCGTACGG<br>CGTAAACCGGGTATGGATCGTTCCGACCTGTTTAACGTTAACGCCGGC<br>ATCGTGAAAAACCTGGTACAGCAAGTTGCGAAAACCTGCCCGAAAGCG<br>TGCATTGGTATTATCACTAACCCGGTTAACACCACAGTTGCAATTGCT<br>GCTGAAGTGCTGAAAAAAGCCGGTGTTTATGACAAAAACAAACTGTTG<br>GGCGTTACCACGCTGGATATCATTCGTTCCAACACCTTTGTTGCGGAA<br>CTGAAAGGCAAACAGCCTGGCGAAGTTGAAGTGCCGGTTATTGGCGGT<br>CACTCTGGTGTTACCATTCTGCCGCTGCTGTCACAGGTTCCTGGCGTT<br>AGTTTTACCGAGCAGGAAGTCGCTGATCTGACCAAACGCATCCAGAAC<br>GCGGGTACTGAAGTGGTTGAAGCGAAGGCCGGTGGCGGGTCTGCAACC<br>CTGTCTATGGGCCAGGCAGCTGCACGTTTTGGTCTGTCTCTGGTTCGT<br>GCACTGCAGGGCGAACAAGGCGTTGTCGAATGTGCCTACGTTGAAGGC<br>GACGGTCAGTACGCCCGTTTCTTCTCTCAACCGCTGCTGCTGGGTAAA<br>AACGGCGTGGAAGAGCGTAAATCTATCGGTACCCTGAGCGCATTTGAA<br>CAGAACGCGCTGGAAGGTATGCTGGATACGCTGAAGAAAGATATCGCC<br>CTGGGGCAAGAGTTCGTTAATAAGTAA |
| fdh_Sn | ATGGCCAAAATTTTATGCGTATTATACGATGATCCGGTCGATGGCTAC<br>CCAAAAACCTATGCACGCGATGACTTGCCGAAGATCGACCACTATCCG<br>GGGGGTCAGACACTGCCGACGCCAAAGGCGATTGACTTCACTCCTGGT<br>GCATTGTTGGGGTCCGTAAGCGGTGAGTTGGGATTACGCAAATACCTT<br>GAGGCTAACGGTCATACGTTTGTCGTGACGAGTGACAAAGACGGGCCG<br>GACTCAGTGTTTGAACGCGAACTGGTGGACGCGGACGTTGTAATTTCA<br>CAGCCTTTCTGGCCGGCTTATTTAACCCCGGAGCGCATTGCAAAAGCC<br>AAGAATCTTAAGCTGGCTTTGACTGCTGGTATTGGTTCGGACCATGTG<br>GATCTGCAGTCTGCAATTGACCGTGGCATCACAGTGGCAGAGGTCACT<br>TATTGTAATTCCATTAGTGTTGCGGAGCATGTAGTAATGATGATTCTG<br>GGGTTAGTTCGCAACTACATCCCTTCCCATGACTGGGCTCGTAAGGGC<br>GGATGGAACATCGCGGATTGCGTCGAGCATTCGTATGACCTTGAGGGT<br>ATGACAGTGGGCTCAGTCGCCGCTGGCCGTATTGGCCTTGCTGTGCTG<br>CGCCGTTTAGCGCCCTTCGATGTCAAATTGCATTATACGGATCGTCAT<br>CGCTTACCAGAAGCCGTCGAGAAGGAACTGGGTTTAGTTTGGCACGAC<br>ACACGCGAGGACATGTACCCACATTGTGATGTCGTCACGTTAAACGTC<br>CCCCTTCATCCTGAAACCGAGCACATGATTAACGACGAAACTTTAAAG<br>CTTTTTAAACGCGGTGCGTACATTGTAAATACGGCTCGCGGGAAGTTG<br>GCTGACCGTGATGCTATTGTACGTGCCATTGAATCCGGACAACTGGCT<br>GGGTATGCTGGGGATGTGTGGTTCCCTCAACCAGCACCAAAGGACCAT<br>CCATGGCGTACCATGAAATGGGAGGGAATGACACCTCACATCAGTGGG |

| | |
|---|---|
| | ACGTCACTTAGCGCGCAAGCCCGTTACGCTGCTGGGACACGCGAAATT<br>TTAGAATGCTTTTTTGAGGGGCGTCCGATTCGCGATGAATACCTGATC<br>GTGCAAGGCGGCGCGTTGGCCGGGACCGGCGCACATTCGTACTCTAAA<br>GGGAACGCTACGGGTGGAAGTGAGGAGGCGGCTAAGTTTAAAAAAGCT<br>GGATAA |
| pyc_Re | ATGCCGATTAGTAAAATTTTGGTTGCAAATCGTAGTGAGATTGCTATC<br>CGTGTCTTTCGCGCGGCTAATGAACTGGGGATTAAGACCGTTGCGATT<br>TGGGCCGAAGAGGACAAGCTGGCGTTGCACCGCTTCAAAGCTGACGAA<br>TCGTACCAGGTTGGTCGCGGTCCTCATCTGGCCCGCGATCTGGGCCCG<br>ATCGAGAGTTATCTGAGCATTGACGAAGTTATCCGCGTAGCTAAGCTG<br>TCCGGTGCTGATGCTATCCACCCCGGCTATGGTCTTTTATCTGAGTCT<br>CCAGAATTTGTGGATGCGTGCAACAAAGCCGGGATCATTTTCATTGGT<br>CCGAAAGCCGACACGATGCGCCAGTTGGGAAACAAGGTCGCGGCCCGC<br>AATCTGGCTATTTCCGTAGGCGTCCCGGTAGTGCCTGCTACAGAACCG<br>CTGCCCGACGATATGGCAGAAGTAGCTAAGATGGCCGCGGCAATTGGG<br>TATCCAGTGATGCTGAAGGCATCATGGGGAGGAGGAGGACGCGGTATG<br>CGCGTAATTCGTTCAGAAGCGGATCTTGCGAAGGAAGTTACAGAGGCC<br>AAACGCGAAGCCATGGCGGCGTTTGGTAAAGACGAGGTGTACCTGGAG<br>AAATTGGTAGAACGCGCACGCCACGTCGAGAGTCAAATTTTAGGCGAT<br>ACCCATGGCAACGTCGTGCACTTATTCGAACGCGATTGCTCAGTGCAA<br>CGTCGTAATCAAAAGGTTGTGGAGCGTGCACCCGCTCCATATCTGTCC<br>GAAGCACAACGTCAAGAACTGGCCGCATACAGTCTGAAAATTGCAGGC<br>GCTACTAACTATATCGGCGCCGGAACCGTGGAATACTTAATGGACGCC<br>GACACGGGGAAGTTCTATTTTATCGAGGTTAACCCTCGTATTCAGGTA<br>GAACACACAGTGACCGAGGTGGTGACGGGGATCGACATCGTTAAAGCC<br>CAGATTCATATTTTAGACGGCGCCGCCATTGGGACTCCCCAGAGTGGG<br>GTCCCTAATCAAGAAGACATTCGTCTGAATGGTCACGCTTTGCAATGC<br>CGCGTAACGACAGAGGATCCAGAACACAACTTCATCCCGGACTATGGC<br>CGCATCACAGCCTATCGCTCGGCCAGTGGTTTCGGTATCCGCTTGGAC<br>GGTGGAACGTCCTACAGTGGCGCCATCATCACCCGTTATTACGACCCA<br>TTGCTGGTCAAAGTCACTGCTTGGGCACCCAACCCATTAGAAGCAATT<br>TCTCGTATGGACCGCGCCTTACGTGAGTTCCGTATCCGCGGTGTTGCG<br>ACGAACCTTACGTTCTTGGAAGCTATCATCGGCCACCCTAAGTTTCGT<br>GACAATAGCTATACCACGCGCTTTATCGACACTACCCCCGAATTATTT<br>CAACAGGTGAAGCGCCAGGACCGCGCCACTAAGTTGCTGACTTATCTT<br>GCAGACGTAACAGTTAATGGGCATCCGGAAGCTAAGGACCGCCCCAAG<br>CCTCTTGAAAACGCCGCGCGCCCCGTGGTACCGTATGCCAATGGCAAC<br>GGTGTCAAGGACGGCACTAAGCAATTACTTGACACATTAGGGCCGAAA<br>AAGTTTGGCGAATGGATGCGTAACGAGAAACGTGTCCTTTTGACAGAT<br>ACCACAATGCGCGACGGCCATCAATCCTTGTTGGCCACCCGCATGCGC<br>ACTTATGATATTGCACGTATCGCTGGGACCTATTCGCACGCCCTTCCC<br>AACTTGTTGAGTCTGGAGTGCTGGGGGGGTGCCACTTTCGATGTGTCG<br>ATGCGCTTCCTTACCGAGGACCCTTGGGAGCGCTTGGCTCTGATTCGT<br>GAGGGGGCGCCAAATCTTCTGCTTCAAATGTTACTGCGTGGCGCCAAT<br>GGGGTAGGATATACGAACTACCCTGACAACGTCGTCAAATACTTCGTC |

| | |
|---|---|
| | CGCCAAGCTGCTAAAGGAGGAATCGATCTGTTTCGCGTTTTCGACTGT<br>TTAAATTGGGTGGAGAATATGCGTGTGTCTATGGATGCGATTGCCGAG<br>GAGAATAAGCTTTGCGAAGCTGCCATTTGTTATACGGGGGATATTCTG<br>AATTCCGCCCGTCCTAAGTATGACTTGAAATACTATACTAACCTTGCG<br>GTCGAATTGGAGAAGGCTGGAGCTCACATTATCGCTGTCAAGGACATG<br>GCAGGATTGCTGAAGCCTGCCGCAGCAAAAGTGCTTTTCAAGGCTCTT<br>CGTGAAGCAACGGGTCTGCCAATTCATTTCCATACCCATGATACCAGC<br>GGAATCGCAGCTGCTACTGTGTTAGCGGCGGTGGAAGCAGGAGTAGAT<br>GCCGTAGACGCAGCTATGGACGCCCTGAGTGGGAACACGTCGCAACCA<br>TGCCTGGGCTCAATCGTAGAAGCCCTTTCAGGTTCAGAACGCGATCCA<br>GGATTGGACCCTGCGTGGATTCGTCGCATTTCTTTTTACTGGGAGGCT<br>GTTCGTAACCAATACGCAGCCTTTGAGAGTGACTTGAAGGGCCCTGCT<br>AGCGAAGTCTACTTGCATGAAATGCCCGGAGGTCAATTTACAAATCTG<br>AAGGAGCAAGCCCGCTCTCTGGGCTTAGAAACTCGCTGGCACCAAGTA<br>GCACAGGCGTACGCGGATGCGAACCAGATGTTCGGCGATATCGTTAAG<br>GTAACGCCGTCGAGCAAAGTCGTTGGCGACATGGCGTTGATGATGGTT<br>TCTCAGGATCTGACTGTGGCGGATGTTGTCAGCCCAGATCGCGAGGTA<br>AGCTTCCCTGAGTCAGTCGTAAGTATGCTTAAAGGAGACCTGGGACAG<br>CCGCCTTCGGGGTGGCCGGAAGCTTTACAAAAGAAAGCTCTGAAAGGC<br>GAGAAACCGTACACTGTACGTCCTGGTAGTTTGCTTAAGGAAGCCGAT<br>CTGGACGCCGAGCGTAAGGTTATCGAGAAGAAATTGGAGCGCGAGGTG<br>AGTGATTTTGAATTCGCAAGTTATTTAATGTATCCAAAGGTGTTTACA<br>GATTTTGCCCTTGCATCAGATACTTATGGTCCTGTATCAGTTTTGCCC<br>ACCCCTGCCTATTTTTATGGTTTAGCCGATGGAGAGGAGCTTTTTGCC<br>GATATCGAAAAAGGCAAGACTCTGGTGATTGTTAATCAGGCAGTAAGC<br>GCCACGGACAGCCAAGGCATGGTTACAGTATTCTTTGAGTTAAACGGC<br>CAGCCCCGTCGTATCAAAGTACCTGACCGCGCTCATGGAGCTACGGGA<br>GCCGCCGTTCGCCGCAAAGCAGAACCGGGGAACGCCGCTCATGTTGGG<br>GCCCCGATGCCGGGGGTAATTAGCCGTGTTTTTGTTTCGTCAGGACAG<br>GCAGTAAATGCCGGGGACGTGCTTGTTTCTATCGAGGCGATGAAAATG<br>GAAACCGCAATCCATGCGGAGAAGGATGGTACGATCGCTGAAGTATTG<br>GTTAAAGCTGGCGATCAAATCGATGCTAAGGACCTGCTGGCGGTGTAT<br>GGGGGTTAA |
| sda_Ec | ATGATCTCGCTGTTCGATATGTTCAAAGTAGGTATCGGACCTAGCAGC<br>AGCCATACCGTCGGGCCGATGAAAGCCGGGAAACAATTTGTCGATGAC<br>TTGGTAGAGAAAGGTTTATTAGACTCCGTAACACGCGTGGCCGTGGAC<br>GTATATGGGTCCCTTTCATTGACAGGAAAGGGGCACCACACCGATATC<br>GCCATCATTATGGGGCTTGCAGGAAACGAACCAGCGACCGTGGATATC<br>GACTCTATTCCGGGGTTTATCCGCGATGTCGAGGAGCGTGAGCGCCTT<br>TTGCTGGCTCAGGGTCGCCACGAGGTTGATTTCCCCCGTGACAATGGT<br>ATGCGTTTCCACAATGGCAACCTTCCGCTTCACGAAAACGGCATGCAG<br>ATTCACGCATACAACGGGGACGAAGTAGTGTACTCTAAGACATATTAC<br>TCCATTGGCGGGGGATTTATTGTTGATGAGGAACATTTCGGACAAGAT<br>GCGGCAAATGAAGTGTCCGTACCTTATCCATTTAAGTCAGCCACGGAA<br>TTGCTTGCCTACTGCAACGAGACTGGATATTCCTTGTCTGGCTTAGCG |

| | |
|---|---|
| | ATGCAGAATGAGCTGGCTTTGCACTCAAAGAAGGAGATTGATGAGTAC<br>TTTGCGCACGTGTGGCAAACCATGCAAGCGTGTATCGATCGCGGAATG<br>AACACAGAAGGGGTCTTACCGGGACCGTTACGTGTCCCACGTCGTGCC<br>TCGGCTCTGCGTCGTATGCTTGTATCATCTGATAAATTATCTAATGAC<br>CCCATGAATGTTATTGACTGGGTGAATATGTTCGCTCTGGCAGTGAAT<br>GAAGAAAATGCGGCCGGGGGACGTGTAGTTACTGCTCCGACAAACGGT<br>GCCTGTGGAATTGTGCCTGCGGTTCTTGCTTACTACGACCATTTCATC<br>GAGTCGGTCTCCCCGGACATCTACACACGTTATTTCATGGCCGCTGGC<br>GCAATTGGTGCCCTTTACAAGATGAACGCGTCGATTTCGGGGGCCGAG<br>GTAGGGTGCCAAGGCGAGGTGGGAGTGGCTTGCTCAATGGCAGCGGCC<br>GGCTTGGCGGAATTGCTGGGAGGAAGTCCAGAGCAGGTCTGCGTCGCG<br>GCCGAAATCGGTATGGAGCACAATCTGGGGTTAACCTGTGATCCGGTC<br>GCGGGGCAGGTCCAAGTGCCTTGTATTGAGCGCAATGCTATTGCGTCT<br>GTGAAAGCTATCAATGCTGCCCGCATGGCGCTGCGTCGTACTTCCGCG<br>CCACGTGTATCTCTGGATAAAGTTATTGAAACCATGTATGAAACCGGG<br>AAGGACATGAACGCCAAGTATCGCGAAACCTCTCGTGGGGGACTTGCA<br>ATCAAAGTGCAGTGCGATTGA |
| sda_Bsub | ATGAAATACCGCAGTGTATTCGATATTATTGGCCCAGTCATGATTGGT<br>CCTTCAAGTTCTCACACTGCTGGTGCAGCCCGTATTGGGCGTGTGGCC<br>CGCTCGTTGTTCGGTCGCGAGCCTGAGCGCATTATTGTTTCGTTTTAT<br>GGATCATTCGCAGAGACGTATAAAGGCCATGGAACCGATGTCGCCATT<br>ATTGGAGGCTTACTTGATTTTGACACGTTTGATGAGCGCATCAAGACG<br>GCGATCCAGATCGCGGAAGCCAAAGGCATCGACATCGAGTTCCGCGTG<br>GAGGATGCGGTCCCGGTACATCCTAACACAGCCAAGATTACGATTTCA<br>GATGAAAAAGGTGAACTTGAACTTACGGGGATTTCGATCGGCGGGGGA<br>AAAATTGAAATCACTGAATTAAACGGATTTGAATTACGCCTTAGCGGG<br>AATCATCCAGCTATCCTTGTAGTGCACAACGATAAATTCGGGACAATT<br>GCTGGAGTCGCTAACGTGCTTGCGAAGTTTAGTATTAATGTAGGACAC<br>ATGGAAGTCGCCCGTAAAGACATTGGTCAACTGGCCCTTATGACGATC<br>GAAGTAGACCAGAACATTGACGACCATATTCTGGACGAACTTTCAAAA<br>TTGCCCAACATTATTCAGGTTACTAAGATTGCAGATTGA |
| sda_Msmeg | ATGGATCTGGTTACCTTAGATGACATTTCGGGTGCAGCTGCCCGTATT<br>GCGGCCGATATCGTCCGTACACCTTTATTGGCGGCCGATTGGGGGGAT<br>CCTCGCTGTCCATTATGGCTTAAAGCTGAAACACTGCAGCCCATCGGC<br>GCATTCAAAATTCGCGGTGCTTTTAACGCTCTGGGTCGCCTTGACACG<br>CACACGCGCGCACGTGGCGTAGTGGCATATTCTTCGGGGAATCATGCC<br>CAGGCGGTAGCATATGCAGCGGCTGCCTATGGTGTTCCCGCCCACATT<br>GTCATGCCAGAGGAGACACCAGCCGTCAAAGTCGAGGCAACTCGTCGT<br>CGTGGAGCCCATGTGGTGCTGTGTGGTGCCGGTGAACGTGAACGCACC<br>GCCGCAGAGTTAGTGGAGAAGACGGGAGCCGTACTGATCCCGCCTTTC<br>GACCACCCCGATATTATCGCTGGTCAGGGGACGATTGGAATTGAGATT<br>GCGGAGGATCTTCCCGAATTGGCTACAGTGTTGATCCCGGTAAGCGGC<br>GGGGGTTTGGCGAGTGGGATTGGAACTGCCATCCGCGCGTTGCGTCCT<br>AAGGCAAAAATTTTCGCAGTAGAGCCGGAGTTAGCGGCGGATACCGCA<br>GAGAGCTTAGCGTTGGGAAGTATTGTCGAGTGGCCAGTTGCCAAGCGC |

| | |
|---|---|
| | AACCGCACAATTGCAGACGGTTTACGTTCCACACCATCAGAGTTAACG<br>TTTGCCCACCTGCGCCAAGTCATTGATGACGTCATTACGGTTTCGGAG<br>GATGAAATCCGTTCAGCAGTACGTGAGCTGGCATTGCGTGCGCGTTTG<br>GTTGCAGAACCGTCCGGAGCGGTATCTCTTGCCGGGTATCGCAAGGCG<br>GCTTTACCTGATGGCAGTGCGGTTGCCATCGTGAGCGGGGGAACATT<br>GAACCGGCACAACTTGCGGCTATCCTGGCTGGCGGATAA |
| sda_Lpneu | ATGAATATTTCTGTCTTCGATTTGTTCAGTATCGGGATTGGCCCTTCT<br>TCATCCCACACTGTCGGCCCAATGTTGGCAGCAAATGCCTTCCTTCAA<br>CTTCTTGAGCAGAAGAACCTTTTCGATAAAACGCAACGCGTAAAAGTG<br>GAGCTGTATGGCAGTTTGGCGCTGACCGGCAAAGGCCATGGAACCGAC<br>AAGGCGATCTTGAACGGTCTGGAAAACAAGGCCCCTGAGACAGTGGAC<br>CCGGCCTCCATGATCCCACGCATGCACGAGATCTTAGATTCCAATTTG<br>CTGAACCTGGCTGGGAAGAAGGAGATTCCCTTTCACGAAGCAACCGAT<br>TTCCTGTTCTTGCAAAAAGAACTGCTTCCAAAGCACTCAAATGGCATG<br>CGCTTCTCGGCCTTCGATGGGAATGCCAACCTGCTTATTGAACAGGTG<br>TATTATAGTATCGGCGGCGGGTTCATTACAACAGAAGAGGATTTTGAT<br>AAGTCCACTACTGATACTAATCCACCTCCTTATCCTTTTGCGACAGCA<br>ACTGAGTTACTTAAATTATGCAAAAAGCATCATCTTACGATCGCTGAA<br>CTGATGCTTGTTAATGAGAAGACTTGGCGTAGTTCTGCTGAAATCCAT<br>AAAGGTATCCTTGACATTGCTAAGGTAATGGACGATTGTATCAACAAT<br>GGTTGCAAACATGATGGTGTCCTGCCGGGAGGCTTAAATTTAAAACGC<br>CGCGCTCCCGACTTGTACCGCAAACTTATTGAGCAGAAGGGAGTGAAA<br>TCTGTATTTGAGCAGTCTGACATCATGAATCACTTAAACTTGTACGCC<br>ATGGCGGTCAATGAGGAAAATGCAGCTGGCGGCCGTATCGTGACCGCA<br>CCTACAAATGGGGCGGCAGGCATCATCCCGGCGGTACTGAAGTATTGC<br>CAACAGGCCCACGACCGCATGTCGAATGAGGACATTTACACCTATTTC<br>TTGACCGCTGCGGCTATTGGCATTCTGTATAAGAAGGGCGCGTCAATT<br>TCCGGAGCGGAAGTAGGTTGTCAGGGTGAAGTAGGAGTAGCATCCTCT<br>ATGGCAGCCGCGGGACTTACTGCAGTTCTGGGAGGGACCATCGAACAA<br>GTGGAGAACGCGGCTGAGATCGCCATGGAGCACCATCTGGGAATGACG<br>TGCGATCCTGTCTTGGGCTTAGTACAGATCCCGTGTATCGAGCGCAAC<br>GCCATGGGTGCCGTTAAGGCTGTTAATGCGACACGCATGGCGTTAATC<br>GGAGATGGGCAGCACCAAATTTCCCTGGATAAGGTGATCAAGACGATG<br>AAGCAGACCGGCATGGACATGCAGTCAATTTACAAGGAAACGTCTATG<br>GGTGGTCTGGCAGTGAACCTTCCAGAGTGCTAA |
| TdcB_Ec | atgcatattacatacgatctgccggttgctattgatgacattattgaa<br>gcgaaacaacgactggctgggcgaatttataaaacaggcatgcctcgc<br>tccaactattttagtgaacgttgcaaaggtgaaatattcctgaagttt<br>gaaaatatgcagcgtacgggttcatttaaaattcgtggcgcatttaat<br>aaattaagttcactgaccgatgcggaaaaacgcaaaggcgtggtggcc<br>tgttctgcgggcaaccatgcgcaaggggtttccctctcctgcgcgatg<br>ctgggtatcgacggtaaagtggtgatgccaaaaggtgcgccaaaatcc<br>aaagtagcggcaacgtgcgactactccgcagaagtcgttctgcatggt<br>gataacttcaacgacactatcgctaaagtgagcgaaattgtcgaaatg<br>gaaggccgtattttttatcccaccttacgatgatccgaaagtgattgct |

| | |
|---|---|
| | ggccagggaacgattggtctggaaattatggaagatctctatgatgtc<br>gataacgtgattgtgccaattggtggtggcggtttaattgctggtatt<br>gcggtggcaattaaatctattaacccgaccattcgtgttattggcgta<br>cagtctgaaaacgttcacggcatggcggcttctttccactccggagaa<br>ataaccacgcaccgaactaccggcaccctggcggatggttgtgatgtc<br>tcccgcccgggtaatttaacttacgaaatcgttcgtgaattagtcgat<br>gacatcgtgctggtcagcgaagacgaaatcagaaacagtatgattgcc<br>ttaattcagcgcaataaagtcgtcaccgaaggcgcaggcgctctggca<br>tgtgctgcattattaagcggtaaattagaccaatatattcaaaacaga<br>aaaaccgtcagtattatttccggcggcaatatcgatctttctcgcgtc<br>tctcaaatcaccggtttcgttgacgcttaa |
| sds_Dd | ATGGGACTGAAGACTACAATCAGCCCTGACTCATCGTTTGATAAAATC<br>ATCAATACTAAGACGAGTCCTCCTCTGCATATCAATAGTCCCATGTTG<br>GAGTCTTTGGCCTTAAGCAAATTATTCAAGGAGGAAAACGCGAAAGTC<br>TGGATGAAGGTAGACGCTCTTCAACCTTCTGGATCGTTTAAGATCCGT<br>GGCGTAGGATTGCTTTGCAACCAGTTGCTGAAGGAAAAGAAATCTAAA<br>AACGAAGAGGCCCACTTCATCTGTTCTTCGGGCGGAAACGCGGGCAAG<br>TCAGTGGCGTACGCAGGACGTAAACTTAACGTTAAGACCACTATCGTG<br>CTTCCAAACACCATTCCGGAGGCGACCATCGAGAAGATTAAAGATGAG<br>GGCGCTAATGTTATTGTGCACGGAACCATCTGGGATGAGGCGAACACC<br>TTTGCGCTTGAGCTTGCTGAGAAAGAGGGCTGCACTGACTGCTACATT<br>CATCCCTTCGATCACCCTCTTCTTTGGGAGGGCCACAGTACCATGATT<br>GATGAGATTTATCAAGATGTCCAGAACGGAGTATGCGAGAAGCCAGAC<br>GTAATTCTGTTCTCAGTGGGTGGCGGCGGCATGATGATCGGCATCTTA<br>CAAGGTTTAGACCGCTACGGATGGAACGATATTCCGATTGTAACCGTG<br>GAGACGGTCGGTAGCCACTCTTTTTGGAAATCCTTCCAAGAAAAACAA<br>CTTACTAAATTAGACGTAAGTGAGGTCACCAGTGTGATCAAAACATTA<br>TCCACGCGTTCAGTTTGCTCAGAAGCATGGGAGATTTCTAAACGCTTT<br>AATATTAAACCCATTCTTGTCACCGACCGTGACGCTGTAGACGCATGC<br>TTAAAGTTCGTTGACGACGAACGCATTCTGGTCGAGCCTAGTTGCGGT<br>GCTACTCTGTCGGTGCTTTACTCGAAGAAATTGTCTACTTTGTTAGAT<br>ATTAACTCTAAAAACATCTTAACCATTGTATGTGGTGGCAACGGGACC<br>AGTATTCTTCAGTTGAACGACTTATTGCAGACGCTGCCGAAGtaa |
| glyA_Ec | atgttaaagcgtgaaatgaacattgccgattatgatgccgaactgtgg<br>caggctatggagcaggaaaaagtacgtcaggaagagcacatcgaactg<br>atcgcctccgaaaactacaccagcccgcgcgtaatgcaggcgcagggt<br>tctcagctgaccaacaaatatgctgaaggttatccgggcaaacgctac<br>tacggcggttgcgagtatgttgatatcgttgaacaactggcgatcgat<br>cgtgcgaaagaactgttcggcgctgactacgctaacgtccagccgcac<br>tccggctcccaggctaactttgcggtctacaccgcgctgctggaacca<br>ggtgataccgttctgggtatgaacctggcgcatggcggtcacctgact<br>cacggttctccggttaacttctccggtaaactgtacaacatcgttcct<br>tacggtatcgatgctaccggtcatatcgactacgccgatctggaaaaa<br>caagccaaagaacacaagccgaaaatgattatcggtggtttctctgca<br>tattccggcgtggtggactgggcgaaaatgcgtgaaatcgctgacagc |

| | |
|---|---|
| | atcggtgcttacctgttcgttgatatggcgcacgttgcgggcctggtt gctgctggcgtctacccgaacccggttcctcatgctcacgttgttact accaccactcacaaaaccctggcgggtccgcgcggcggcctgatcctg gcgaaaggtggtagcgaagagctgtacaaaaaactgaactctgccgtt ttccctggtggtcagggcggtccgttgatgcacgtaatcgccggtaaa gcggttgctctgaaagaagcgatggagcctgagttcaaaacttaccag cagcaggtcgctaaaaacgctaaagcgatggtagaagtgttcctcgag cgcggctacaaagtggtttccggcggcactgataaccacctgttcctg gttgatctggttgataaaaacctgaccggtaaagaagcagacgccgct ctgggccgtgctaacatcaccgtcaacaaaaacagcgtaccgaacgat ccgaagagcccgtttgtgacctccggtattcgtgtaggtactccggcg attacccgtcgcggctttaaagaagccgaagcgaaagaactggctggc tggatgtgtgacgtgctggacagcatcaatgatgaagccgttatcgag cgcatcaaaggtaaagttctcgacatctgcgcacgttacccggtttac gca |
| MtdA_Mex | atgtctaagaaactgctctttcagtttgacactgatgcaactccgtct gtatttgacgttgttgttggctatgacggcggtgcagaccatattact ggctatggcaatgttactcccgacaatgttggcgcatatgttgacggc actatttatactcgtggaggcaaagagaaacagtctacagcaatcttt gttggcggcggcgacatggcagcaggcgagcgggtatttgaggcagta aagaagcgtttctttggcccgtttcgcgtttcttgtatgctggattct aatggctctaatactactgcagcagcaggcgttgcactcgttgttaaa gcagcaggcggctctgttaaaggcaagaaagcagttgttctcgcaggt actggtccggttggtatgcgctctgcagctctgttagccggcgagggc gcagaggttgttctgtgtgggcgcaaactcgacaaagcacaggcagca gcagattctgttaataaacgcttcaaagttaatgttactgcagcagag actgcagacgacgcatctcgcgcagaggccgtgaaaggcgcacatttt gtctttactgcaggtgcaattggccttgaactgctgccgcaggcagca tggcagaatgagtcttctattgaaattgtggccgattataatgcacag ccgccgctcggcattggcgggattgatgcaactgacaaaggcaaagaa tatggcggaaaacgcgcatttggtgcgctcggcattggcggcttgaaa ctcaaactgcatcgcgcatgtattgcaaaactgtttgagtcttctgaa ggtgtatttgatgcagaggagatttataaactggcaaaagaaatggca tga |
| Fch_Mex | atggctggcaatgagactattgaaacattcttggacggcctggcatca tctgctccgactcccggcggcggcggtgcagcagcaatttctggcgca atgggcgcagcacttgtttctatggtttgcaatcttactattggcaag aagaaatatgttgaggttgaggcagacttaaaacaggttctggagaaa tctgaaggcctgcgccgcactctcactggcatgattgcagacgacgtt gaagcctttgacgcagttatgggcgcttatgggctgccgaagaatact gacgaagagaaagcagcacgcgcagcaaagattcaagaggcactcaaa actgcaactgacgttccgctcgcatgttgtcgcgtttgtcgcgaggtt attgatctggcagagattgttgcagagaaaggcaatctcaatgttatt tctgatgcaggcgttgcagtgctctctgcttatgcaggtctgcgctct gctgcacttaatgtctatgtaaatgcaaaaggcctcgacgaccgcgca |

| | |
|---|---|
| | tttgcagaggagcggcttaaagagctggagggcctactggctgaggca ggtgcactcaatgagcgaatttatgagactgttaaatctaaagtgaat tga |
| Ftl_Mex | atgccgagcgatattgaaattgcacgcgctgctactctgaaaccgatt gcgcaagttgcggagaaactgggtattccggacgaggctcttcataat tatggcaaacatatcgctaaaatcgaccatgactttattgcttctctt gagggtaaaccagagggcaaacttgttctggttactgctatttcgccg actccagctggcgagggcaaaactactactactgttggtctgggcgat gctctcaaccgcattggcaaacgtgctgttatgtgtctgcgcgagccc tctctcggcccctgttttggcatgaaaggcggcgctgctggtggcggc aaagctcaggttgttccgatggagcagattaatctgcacttcaccggc gattttcacgctattacttctgctcactctctcgctgctgctctgatt gataaccatatttattgggctaacgaactgaatattgacgttcgccgc attcattggcgccgcgttgttgatatgaacgatcgggctctgcgcgct attaatcagtctctcggcggcgttgctaatggctttccgcgcgaggat gggtttgacattactgttgcttctgaggttatggctgtgttttgcctc gccaagaatctggctgatcttgaggagcggctcggccgcattgttatt gcagaaactcgcgatcgcaaaccggttactctggctgatgttaaagct actggcgctatgactgttctgctcaaggatgctcttcagccgaatctc gtgcagactctggagggcaacccggctctgattcacggcggcccgttt gctaacattgctcatggctgtaactcggttattgctactcgcactggc ctgcggctcgctgactatactgttactgaggctggctttggcgctgat ctcggcgctgagaaattcattgatattaaatgtcgccagactggcctc aagccctctgctgttgttattgttgctacgattcgcgctctcaaaatg catggcggcgttaacaagaaagatctccaggctgagaatctggatgcg ctggagaaaggttttgcaaatcttgagcgccatgttcacaatgttcgc tcttttggcctgccggttgttgttggtgttaaccacttctttcaggat actgatgctgagcatgttcggttgaaagaactgtgccgcgatcggctt caggttgaggctattacttgtaagcattgggctgagggcggcgcaggc gcagaagcactggcacaggcagttgttaaactggctgaaggcgagcag aaaccgctgacttttgcatatgagaccgaaactaagattactgacaag attaaggcaattgctactaaactgtatggtgctgctgatattcagatt gagtctaaagccgccactaagctcgctggcttcgagaaagatggctat ggtaagctgccggtctgtatggccaagactcaatattcatttctact gatccgactcttatgggcgctccctctggtcatctggtttctgtgcgc gatgttcgcctctctgctggcgctggcttcgttgttgttatttgtggt gagattatgaccatgccgggtctgccgaaggttccagcagcagatact attcgcctcgatgctaacggtcagattgatgggctgttctag |
| ppk12 | ATGATTAACATCTACAAGATCGACAAACTTAACAACTTTAACCTGAAT AACCACAAGACCGACGACTACTCTCTTTGTAAGGATAAGGATACTGCT CTGGAGCTTACTCAGAAGAACATCCAGAAAATTTATGATTATCAGCAA AAATTGTACGCCGAAAAAAAAGAAGGATTGATCATTGCATTCCAAGCA ATGGATGCCGCCGGTAAGGATGGAACTATCCGTGAGGTGTTAAAAGCC CTGGCTCCCCAAGGAGTGCACGAGAAGCCCTTCAAGTCCCCATCGAGT ACCGAGCTTGCCCACGATTACTTGTGGCGCGTACACAATGCCGTACCT |

| | |
|---|---|
| | GAAAAAGGTGAGATCACAATTTTTAACCGTAGTCATTACGAAGACGTA<br>TTGATTGGAAAGGTTAAGGAGCTTTATAAATTCCAAAACAAAGCGGAT<br>CGCATTGACGAGAACACGGTAGTTGACAATCGTTATGAGGATATTCGT<br>AACTTTGAGAAATACTTATATAATAATTCAGTCCGCATTATCAAAATT<br>TTTTTGAACGTGTCTAAAAAAGAACAGGCCGAACGTTTCTTATCGCGT<br>ATTGAGGAACCGGAAAAGAACTGGAAATTTTCCGACAGTGATTTCGAG<br>GAGCGTGTATATTGGGACAAATATCAACAGGCTTTCGAAGATGCAATT<br>AACGCAACGTCTACGAAGGATTGTCCTTGGTACGTCGTGCCTGCTGAT<br>CGCAAATGGTATATGCGCTATGTTGTTTCCGAAATCGTGGTCAAGACC<br>TTAGAAGAGATGAACCCTAAGTACCCTACCGTCACCAAAGAGACCCTG<br>GAGCGTTTTGAAGGTTATCGTACAAAGCTTCTGGAAGAGTATAACTAC<br>GACTTGGATACTATCCGTCCCATCGAGAAA |
| fdh_At | atgaaacaggcgagcagtggagacagtaaaaagatcgtgggagtgttc<br>tataaggccaacgagtacgccaccaagaacccgaacttcctgggctgt<br>gtagagaatgcgctgggcattcgggactggctggaatctcaaggtcac<br>cagtacatcgtgacggacgataaggagggaccagattgcgagctggag<br>aagcacattccagatttacacgtgttaatttcaacgccgttccatcct<br>gcgtatgtcacggccgagcgcattaagaaggcaaaaaacttaaagctg<br>ctgctgacggcgggcatcggaagtgaccacattgacctgcaggcggcg<br>gcagcggcaggtttaacggtggccgaagtgactggaagcaatgtggta<br>agtgtcgcagaggatgaactgatgcggatactgatcttaatgcgcaat<br>ttcgtccctggttataaccaagttgtcaaaggcgagtggaatgtggcc<br>ggcatagcataccgcgcgtatgacctggagggtaaaacgatcggcact<br>gtcggagctggcagaattggcaagttgttattacaaagactgaagccc<br>tttggatgtaacctgctgtaccacgatcgccttcagatggcaccagaa<br>ctggaaaaggagactggagccaagtttgtggaagacttgaatgaaatg<br>cttccgaagtgtgacgttatcgtgatcaatatgccattaacggaaaag<br>acgcgtggaatgttcaataaggagttaattggaaaacttaagaagggt<br>gtgttaattgtgaacaatgcccgtggtgcgattatggagcgccaagcc<br>gttgtggacgctgtcgagtcaggtcatatcggaggttactcaggcgat<br>gtctgggacccgcaacctgctccgaaggatcacccgtggcgatatatg<br>ccaaatcaagcgatgactccgcacacgagtggaacgacaattgacgcg<br>caattaagatatgctgctgggacaaaggatatgctagaacgctatttt<br>aagggcgaggactttcctactgaaaactacatcgttaaggatggcgag<br>ttagcgccacaataccgatga |
| fdh_Cm | ATGAAGATTGTACTTGTCTTGTATGACGCAGGTAAGCACGCAGCGGAC<br>GAGGAGAAGTTGTATGGTTGCACCGAAAACAAGTTAGGGATTGCAAAC<br>TGGCTGAAGGATCAGGGACATGAACTGATTACGACTAGCGACAAAGAA<br>GGTGAAACCAGCGAGCTTGATAAACATATCCCCGACGCGGATATTATC<br>ATTACGACGCCATTCCATCCTGCCTATATCACGAAAGAGCGTCTTGAT<br>AAGGCGAAGAATCTTAAGAGTGTCGTAGTCGCAGGCGTCGGTTCGGAC<br>CACATCGACCTTGATTACATCAACCAAACAGGGAAAAAGATTTCGGTA<br>CTTGAGGTGACAGGTTCAAATGTTGTCTCTGTAGCTGAGCATGTCGTA<br>ATGACGATGTTAGTGCTGGTGCGCAACTTCGTGCCCGCTCATGAACAA<br>ATTATCAACCACGATTGGGAGGTTGCAGCGATTGCCAAGGACGCCTAT |

| | |
|---|---|
| | GATATTGAGGGTAAGACCATCGCTACGATCGGTGCCGGTCGTATCGGC<br>TACCGCGTTCTTGAGCGTTTACTGCCATTCAATCCCAAGGAGTTATTA<br>TATTACTCTCTTCAAGCTCTTCCTAAAGAAGCAGAGGAGAAAGTGGGG<br>GCACGTCGTGTCGAAAATATTGAAGAGTTGGTCGCTCAGGCTGACATC<br>GTCACCGTGAACGCGCCATTACACGCCGGCACAAAGGGTCTTATTAAC<br>AAGGAGCTGCTGAGCAAGTTTAAGAAAGGTGCGTGGCTGGTAAATACG<br>GCACGTGGGGCTATTTGTGTCGCGGAAGACGTTGCCGCAGCTCTGGAG<br>TCAGGTCAGTTGCGTGGATATGGCGGGGATGTATGGTTCCCTCAACCA<br>GCGCCTAAGGATCACCCTTGGCGTGATATGCGTAACAAATACGGCGCA<br>GGTAACGCAATGACTCCGCACTACTCAGGGACAACACTTGACGCACAG<br>ACACGTTATGCGGAGGGAACCAAAAACATTTTGGAAAGCTTTTTCACG<br>GGTAAATTTGATTACCGCCCCCAAGACATCATCCTTTTGAATGGCGAA<br>TACGTTACTAAGGCATACGGAAAGCATGACAAGAAATGA |
| deGFP | AGCTTTTCACTGGCGTTGTTCCCATCCTGGTCGAGCTGGACGGCGACG<br>TAAACGGCCACAAGTTCAGCGTGTCCGGCGAGGGCGAGGGCGATGCCA<br>CCTACGGCAAGCTGACCCTGAAGTTCATCTGCACCACCGGCAAGCTGC<br>CCGTGCCCTGGCCCACCCTCGTGACCACCCTGACCTACGGCGTGCAGT<br>GCTTCAGCCGCTACCCCGACCACATGAAGCAGCACGACTTCTTCAAGT<br>CCGCCATGCCCGAAGGCTACGTCCAGGAGCGCACCATCTTCTTCAAGG<br>ACGACGGCAACTACAAGACCCGCGCCGAGGTGAAGTTCGAGGGCGACA<br>CCCTGGTGAACCGCATCGAGCTGAAGGGCATCGACTTCAAGGAGGACG<br>GCAACATCCTGGGGCACAAGCTGGAGTACAACTACAACAGCCACAACG<br>TCTATATCATGGCCGACAAGCAGAAGAACGGCATCAAGGTGAACTTCA<br>AGATCCGCCACAACATCGAGGACGGCAGCGTGCAGCTCGCCGACCACT<br>ACCAGCAGAACACCCCCATCGGCGACGGCCCCGTGCTGCTGCCCGACA<br>ACCACTACCTGAGCACCCAGTCCGCCCTGAGCAAAGACCCCAACGAGA<br>AGCGCGATCACATGGTCCTGCTGGAGTTCGTGACCGCCGCCGGGATC |

### Supplementary Methods

Methods S1: Comparative sequence analysis of enzyme isoforms

BRENDA Enzyme Database and UniProt were searched for PLP-dependent L-serine ammonia-lyase (EC 4.3.1.17) and threonine ammonia-lyase (EC 4.3.1.19) isoforms from single-celled organisms. Isoforms were down-selected based on previous examples of functional enzyme expression in *E. coli*. Multiple sequence analysis (MSA) to determine the presence of AMP binding sites was conducted using Clustal Omega. Protein sequences were gathered from UniProt. Analysis of homology across organisms was performed manually.

Methods S2: Technoeconomic Analysis

The analysis was carried out assuming electrocatalytically generated formate from $CO_2$ as the sole feedstock and with *in situ* regeneration of cofactors. In our TEA model, we also use the "nth" plant assumption, in which the cost assumptions reflect a future time when the technology is well established, and several plants have already been built and are operating. This simplification allows us to avoid accounting for start-up costs and focus on the cost of operation. The specific "nth" cost and financial assumptions are listed elsewhere [4]. Additional technical assumptions to perform the TEA are listed in **Table S2**. A process block flow diagram of the CFE to produce malate is shown in **Figure 7A**. The process economic results are presented in terms of minimum selling price (MSP) on a dollar per kg basis, which is determined by using a discounted cash flow analysis method. All costs in this article are presented on a 2020 constant US dollar basis.

Malate has a market price of $1.8/kg in 2016 [5]. Its U.S. consumption is 15,100 metric tons/year and 16,900 metric tons/year in 2016 and 2021, respectively, with an annual growth rate of 2-3% [5]. Based on the average ethanol plant scale in the U.S., electrochemical conversion efficiency, and product titers, malate production from the cell free process is estimated at 36,510 metric tons/year. Therefore, the effluent $CO_2$ stream from a single ethanol plant could already fulfill the U.S. malate market. TEA suggests that the MSP of malic acid from this technology is about $9.63/kg **(Table S3)**.
